# Intestinal inflammation modulates the expression of ACE2 and TMPRSS2 and potentially overlaps with the pathogenesis of SARS-CoV-2 related disease

**DOI:** 10.1101/2020.05.21.109124

**Authors:** Mayte Suárez-Fariñas, Minami Tokuyama, Gabrielle Wei, Ruiqi Huang, Alexandra Livanos, Divya Jha, Anais Levescot, Haritz Irizar, Roman Kosoy, Sascha Cording, Wenhui Wang, Bojan Losic, Ryan Ungaro, Antonio Di’Narzo, Gustavo Martinez-Delgado, Maria Suprun, Michael J. Corley, Aleksandar Stojmirovic, Sander M. Houten, Lauren Peters, Mark Curran, Carrie Brodmerkel, Jacqueline Perrigoue, Joshua R. Friedman, Ke Hao, Eric E. Schadt, Jun Zhu, Huaibin M. Ko, Judy Cho, Marla C. Dubinsky, Bruce E. Sands, Lishomwa Ndhlovu, Nadine Cerf-Bensusan, Andrew Kasarskis, Jean Frederic Colombel, Noam Harpaz, Carmen Argmann, Saurabh Mehandru

**Author notes:** Contributed equally and address correspondence to and. Contributed equally (Mayte Suárez-Fariñas, Minami Tokuyama). Author contributions: SM, CA, MSF, MT, GW and DJ drafted the manuscript. SM, CA and MSF designed the study, supervised experimental data collection and coordinated integration of collaboration between all participating laboratories. MT performed immunostaining experiments. GW performed network analysis experiments and RH performed statistical analyses. WW, AD, KH, JZ, BL were involved in various aspects of data generation, curation or analysis. All authors critically reviewed and edited the final version of the manuscript.

## Abstract

The presence of gastrointestinal symptoms and high levels of viral RNA in the stool suggest active Severe Acute Respiratory Syndrome Coronavirus 2 (SARS-CoV-2) replication within enterocytes. Here, in multiple, large cohorts of patients with inflammatory bowel disease (IBD), we have studied the intersections between Coronavirus Disease 2019 (COVID-19), intestinal inflammation and IBD treatment. A striking expression of ACE2 on the small bowel enterocyte brush border supports intestinal infectivity by SARS-CoV-2. Commonly used IBD medications, both biologic and non-biologic, do not significantly impact ACE2 and TMPRSS2 receptor expression in the uninflamed intestines. Additionally, we have defined molecular responses to COVID-19 infection that are also enriched in IBD, pointing to shared molecular networks between COVID-19 and IBD. These data generate a novel appreciation of the confluence of COVID-19- and IBD-associated inflammation and provide mechanistic insights supporting further investigation of specific IBD drugs in the treatment of COVID-19.

## Introduction

Severe acute respiratory syndrome coronavirus 2 (SARS-CoV-2) and the ensuing coronavirus disease 2019 (COVID-19)^1^, have evolved into a global pandemic of unprecedented proportions^2^.

Angiotensin converting enzyme-2 (ACE2) is a carboxypeptidase that catalyzes the conversion of angiotensin I into angiotensin 1-9, and angiotensin II into angiotensin 1-7^3-6^. ACE2 can also be cleaved by serine proteases such as transmembrane serine protease (TMPRSS) 2, TMPRSS11D and TMPRSS1. Early events in the pathogenesis of SARS-CoV-2 infection include attachment of the receptor binding domain of the viral spike (S) protein to epithelial ACE2^7-10^. The S protein is then cleaved by TMPRSS2, which facilitates viral entry into the cytoplasm of the host cell^11^. Following infection with SARS-CoV, ACE2 is downregulated in the lungs, resulting in unopposed renin-angiotensin-aldosterone system (RAAS) and contributing to disease severity^12, 13^.

Inflammatory bowel diseases (IBD) encompassing Crohn’s disease (CD) and ulcerative colitis (UC) are chronic, inflammatory disorders of the gastrointestinal (GI) tract that are treated with conventional immunosuppressive drugs such as corticosteroids, biologic therapies and immunomodulatory drugs^14, 15^. Given that SARS-CoV-2 co-opts receptors expressed by intestinal epithelial cells, COVID-19 has the potential to intersect with the pathogenesis of IBD and by extension, its treatment, at a number of points^16, 17^. For example, ACE2 expression may be potentially be altered during gut inflammation or by IBD medications. Further, immunomodulatory drugs used in IBD therapeutics^14, 15^ could potentially be used in COVID-19 patients to manage the “cytokine storm” associated with severe disease.

Therefore, in this study we systematically examined potential areas of intersection between the uninflamed and inflamed GI tract and COVID-19 disease. The results of this study may improve our molecular understanding of how COVID-19 intersects IBD and may provide a rationale for further investigation of drugs used in IBD therapeutics for use in patients with COVID-19.

## Methods

### Immunofluorescence microscopy

Specimens were obtained via clinical endoscopy during routine care (Table S1, S2). Tissue was formalin fixed and paraffin embedded by the clinical pathology core at our institution. Primary antibodies used included ACE2 (abcam-ab15348, 1:1000), EPCAM (abcam-ab228023, prediluted) and mouse anti-TMPRSS2 (Millipore-MABF2158, 1:500) and staining was performed as detailed in supplementary methods.

## Study cohorts

### 1. Cross sectional cohorts

#### i. The Mount Sinai Crohn’s and Colitis Registry (MSCCR)

Peripheral blood and biopsy whole transcriptome sequencing data was obtained from a cross sectional cohort (∼1200 patients) who were enrolled in the Mount Sinai Crohn’s and Colitis Registry (MSCCR) between December 2013 and September 2016 via a protocol approved by the Icahn School of Medicine at Mount Sinai Institutional Review Board. Analysis of the MSCCR cohort is detailed in supplementary methods.

#### ii. The RISK cohort^18^

ACE2 and TMPRSS2 in treatment-free pediatric CD (<17 years of age) patients was studied using RNA-seq expression profiles from GSE57945, which includes ileal biopsies from endoscopically defined inflamed samples (n=160), non-inflamed (n=53) and non-IBD controls (n=42).

### 2. Longitudinal Cohorts

i. The GSE100833 series which includes expression profiles from the gut of 80 anti-TNFα refractory CD patients and the blood from 226 patients enrolled in a phase 2b crossover trial (CERTIFI trial) with ustekinumab as described previously^19^ (detailed in supplementary methods).
ii. The GSE73661 series which includes gene expression profiles (Affymetrix Human Gene 1.0 ST arrays) from colonic biopsies from moderate-to-severe UC patients enrolled in two Vedolizumab efficacy trials (GEMINI-I and GEMINI LTS)^20^ as detailed in supplementary methods. The GSE73661 series also included 12 non-IBD colonic biopsies and colonic biopsies from 23 UC patients before and 4-6 weeks after first infliximab treatment. Response was defined as endoscopic mucosal healing.

### MSCCR Bayesian gene regulatory network (BGRN) generation

BGRNs can capture fundamental properties of complex systems in states that give rise to complex (diseased) phenotypes^19^. BGRNs were generated from RNA sequence data generated on intestinal biopsies from the MSCCR cohort using their intestinal expression QTL information (eQTLs) as priors. The BGRNs were region-(ileum or colon/rectum) and disease-(CD, UC, and control) specific and included both inflamed and uninflamed biopsies and were constructed using RIMBAnet software^19^ and visualized using Cytoscape 3.7^21^. We also used two publicly available BGRNs from the RISK and the CERTIFI cohort^19^ (supplementary methods).

### Bayesian gene regulatory subnetwork generation

*ACE2 and TMPRSS2 subnetworks*: Gene-centric subnetworks were generated by selecting either ACE2 or TMPRSS2 from various BGRNs and expanding out three to five layers (undirected) to obtain the nearest ACE2 or TMPRSS2 neighbors. The connected subnetworks obtained were generally between 200-500 genes in total.

### IBD Inflammation, COVID-19 and IBD Drug Response-Subnetwork generation

We curated RNA-seq based molecular signatures related to IBD and COVID-19 response by identifying differentially expressed genes (DEGs) as detailed in supplementary methods. Genes found differentially expressed in blood^48^, lung NHBE/A549^31^ or human small intestinal organoids^34^ (hSIO) following SARS-CoV-2 infection; IBD inflammation; or response to medications were separately projected onto various BGRNs allowing for 1 or 2 nearest neighbors depending on the signature sizes. The most connected subnetworks were then extracted to generate model-specific SARS-CoV-2 infection-; IBD inflammation-; or drug-response-associated subnetworks (supplementary methods).

### Pathway and geneset enrichment analysis of subnetworks

Gene subnetworks were tested for functional enrichment using a Fisher’s exact test with Benjamini-Hochberg (BH) multiple test correction on a collection of genesets. The collection of genesets included i) Reactome pathways sourced from Enrichr^22^, i) gene sets from Smillie et al^23^, ii) Huang et al^24^, iii) various macrophage perturbations (e.g. cytokines)^25^, iv) ACE2 co-expressed genes^26^ and v) reported IBD GWAS genes (see supplementary methods). Pathway and geneset enrichment, as well as intersection between networks were tested using a Fisher’s exact test and p-values were adjusted for multiple hypothesis with BH correction.

### Key driver gene analysis

Key driver analysis (KDA) identifies key or “master” driver genes for a given gene set in a given BGRN. We used a previously described KDA algorithm^37^ which is summarized in supplementary methods. Genesets for KDA included those associated with NHBE-COVID-19 infection or IBD inflammation. Key driver genes (KDGs) were summarized by frequency across the networks.

### Geneset variation analysis of SARS-CoV-2-infection gene expression signatures

Expression of COVID-19 response gene signatures derived from whole blood or epithelial models was evaluated in the context of IBD related inflammation were studied using gene set variation analysis (GSVA). For each COVID-19 response signature, a sample-wise enrichment score was quantified from each transcriptomic profile in the MSCCR and CERTIFI cohorts using GSVA. COVID-19 response GSVA scores were then modeled to test the association with patient-derived phenotypic information.

## Results

### Healthy gut segments express ACE2 and TMPRSS2 proteins

To define the localization and distribution of ACE2, immunofluorescence (IF) microscopy was performed on histologically normal GI tissue in 20 adults (9 males, 11 females) and 11 children (7 males, 4 females) (**Figure 1 and Table S1**). ACE2 expression was observed on the small intestinal surface epithelium in all subjects in a continuous distribution with the exception of occasional breaks representing the mucin from goblet cells (Figure 1a-b). In all examined small intestinal segments, ACE2 could be detected on the crypt epithelium, though to a lesser extent than on the surface epithelium. TMPRSS2 expression was less abundant in the small bowel, and when detectable, was exclusively found on crypt epithelium (**Figure S1a-b**). There was no observable age or sex dependence of ACE2 or TMPRSS2 protein expression in the small bowel.

**Figure 1.**
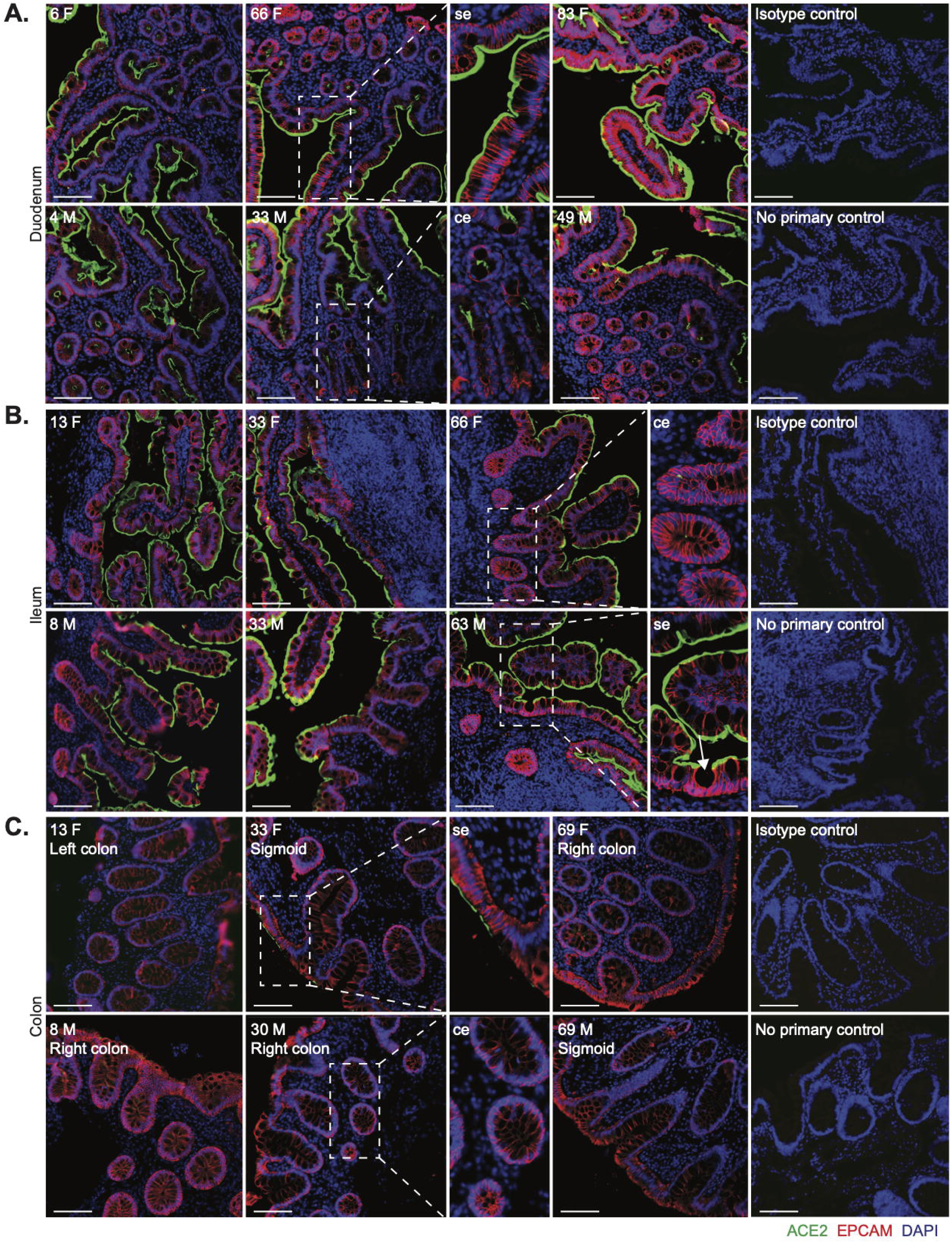
Robust ACE2 expression is found on small bowel surface epithelium in both children and adults. Representative immunofluorescence images of ACE2 (green) and EPCAM (red) counterstained with DAPI (blue) in intestinal biopsies of healthy patients, including magnified images of surface epithelium (se) and crypt epithelium (ce). (**A)** Duodenal biopsies, (**B)** Terminal ileum biopsies (Goblet cell indicated with arrow and (**C)** Colonic biopsies from indicated sites. Patient age (years) and sex (M, male; F, female) as indicated. Isotype controls and no primary controls for each segment are included on the far right of each panel. Scale bar, 100µm.

In the colon, ACE2 expression was patchy and could not be identified in every subject, in contrast to the small bowel (**Figure 1c**). This inconsistency across different donors could not be readily associated with age, sex, ACE inhibitor treatment, nor the colonic segment being examined. In contrast, TMPRSS2 expression was more robust in the colon and was readily detectable on both surface and crypt epithelia (**Figure S1c**). Thus, ACE2 in the healthy gut is higher in the small bowel than the large bowel and inversely, expresses more TMPRSS2 protein in the colon compared to the small bowel.

### ACE2 and TMPRSS2 mRNA expression varies in healthy or inflamed gut segments by region

Next, we examined ACE2 and TMPRSS2 mRNA levels in the intestine of non-IBD controls and IBD patients with active and inactive disease enrolled in MSCCR (**Table S3**). As the number of samples for the colon non-rectum locations was low (**Figure S3a/b**) and no discernable differences were observed, colon non-rectum biopsies were grouped together to increase statistical power. Consistent with the protein data, ACE2 gene expression was higher in the uninflamed ileum compared to the uninflamed colon or rectum. With inflammation, ileal ACE2 mRNA expression was significantly decreased compared to either uninflamed biopsies from IBD patients or non-IBD controls. In contrast, in the rectum ACE2 mRNA expression was increased with inflammation when compared to the uninflamed IBD patients or non-IBD controls (**Figure 2a, upper panels**). There were no significant differences by disease location or between patients with UC versus CD (**Figure S4**). We further validated these results using the pediatric IBD RISK cohort^18^ where ACE2 mRNA was significantly decreased in the ileum of patients with active IBD as well (**Figure 2b**).

**Figure 2:**
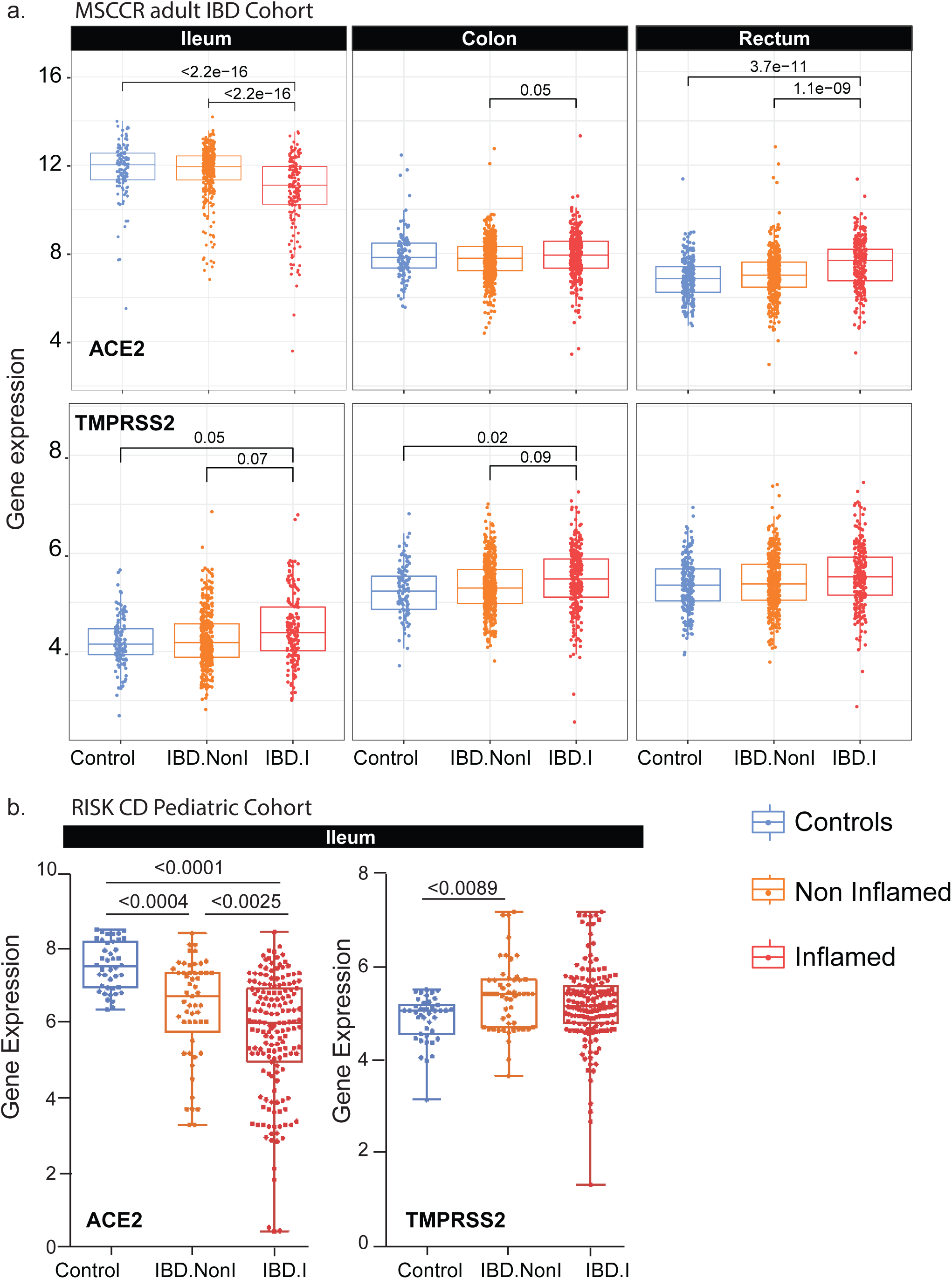
Expression of ACE2 and TMPRSS2 in intestinal biopsies of adult and pediatric IBD patients and controls. **(A)** Box plot summarizing the normalized expression level of ACE2 (upper panel) and TMPRSS2 (lower panel) as measured in ileum or colonic and rectal biopsies from control and IBD MSCCR patients, which were either endoscopically inflamed (IBD.I) or non-inflamed (Non.I). **(B)** Box plot summarizing the normalized expression level of ACE2 (left panel) and TMPRSS2 (right panel) as measured in ileum CD samples from the RISK pediatric cohort. Clinical characteristics of MSCCR IBD patients and biopsies are summarized in Table S3 and S4.

The expression of TMPRSS2 in the MSCCR cohort was moderately higher in the colon compared to ileum. In both ileum and colon biopsies TMPRSS2 expression was found significantly increased in the inflamed relative to non-inflamed samples, although the effect sizes were small (**Figure 2a, lower panels**).

With IF microscopy (**Table S2**), we could not appreciate differences in ACE2 expression in the ileum, which possibly stemmed from the elevated physiological expression of ACE2. In the colon, patchy epithelial ACE2 expression from control non-IBD controls increased in IBD patients with inflammation and this increase was mostly evident on the crypt epithelium (**Figure S2a**).

In the ileum, low intensity TMPRSS2 expression on the crypt epithelium was also comparable in inflamed or uninflamed mucosal segments from IBD patients and in non-inflamed mucosal segments from non-IBD controls. In the colon, TMPRSS2 expression appeared comparable in IBD patients and non-IBD controls. In the rectum, TMPRSS2 expression was enhanced by inflammation (**Figure S2a**).

### Age and gender but not smoking increases ACE2 mRNA in the colon

ACE2 mRNA was higher with age in the uninflamed rectum samples. However, these effects were essentially nullified in the presence of inflammation. A positive association with age in inflamed CD ileum and a negative association in inflamed UC rectum biopsies was observed. ACE2 mRNA in the uninflamed rectum was significantly lower in males versus females but no gender associations in TMPRSS2 mRNA levels were found (**Figure S4b-c**). The expression of ACE2 and TMPRSS2 was similar when comparing active smokers to non-smokers, either between healthy controls or IBD patients (data not shown) and no significant interactions with inflammation status, region or other covariates were found. Thus, age and gender, but not smoking modulates ACE2 but not TMPRSS2 mRNA expression in the IBD colon.

### Non-biological medications: corticosteroids, thiopurines and 5-aminosalicylates reduce ACE2 and TMPRSS2 gene expression in the inflamed colon and rectum but not in the ileum

We further evaluated the impact of non-biologic and biologic medication use (self-reported) on the expression of ACE2 and TMPRSS2 mRNA (Table S3) by propensity matching the MSCCR cohort (see supplementary methods).

In the *ileum* of IBD (CD) patients, corticosteroid, thiopurine or 5-aminosalicylate use had no impact on ACE2 mRNA expression in either inflamed or uninflamed biopsies (**Figure 3a-c**). In the *rectum*, however, a significant decrease in ACE2 mRNA expression was observed with corticosteroid use in inflamed biopsies. A similar decrease of ACE2 mRNA was noticed in thiopurine-treated non-inflamed samples from the rectum. The use of corticosteroids, thiopurine or 5-aminosalicylate did not significantly affect TMPRSS2 mRNA expression in ileum samples. However, each of these three medications significantly decreased TMPRSS2 mRNA expression in inflamed rectum or colon samples. Thus, corticosteroid, thiopurine or 5-aminosalicylate attenuated ACE2 and TMPRSS2 mRNA expression in inflamed colon and rectum.

**Figure 3:**
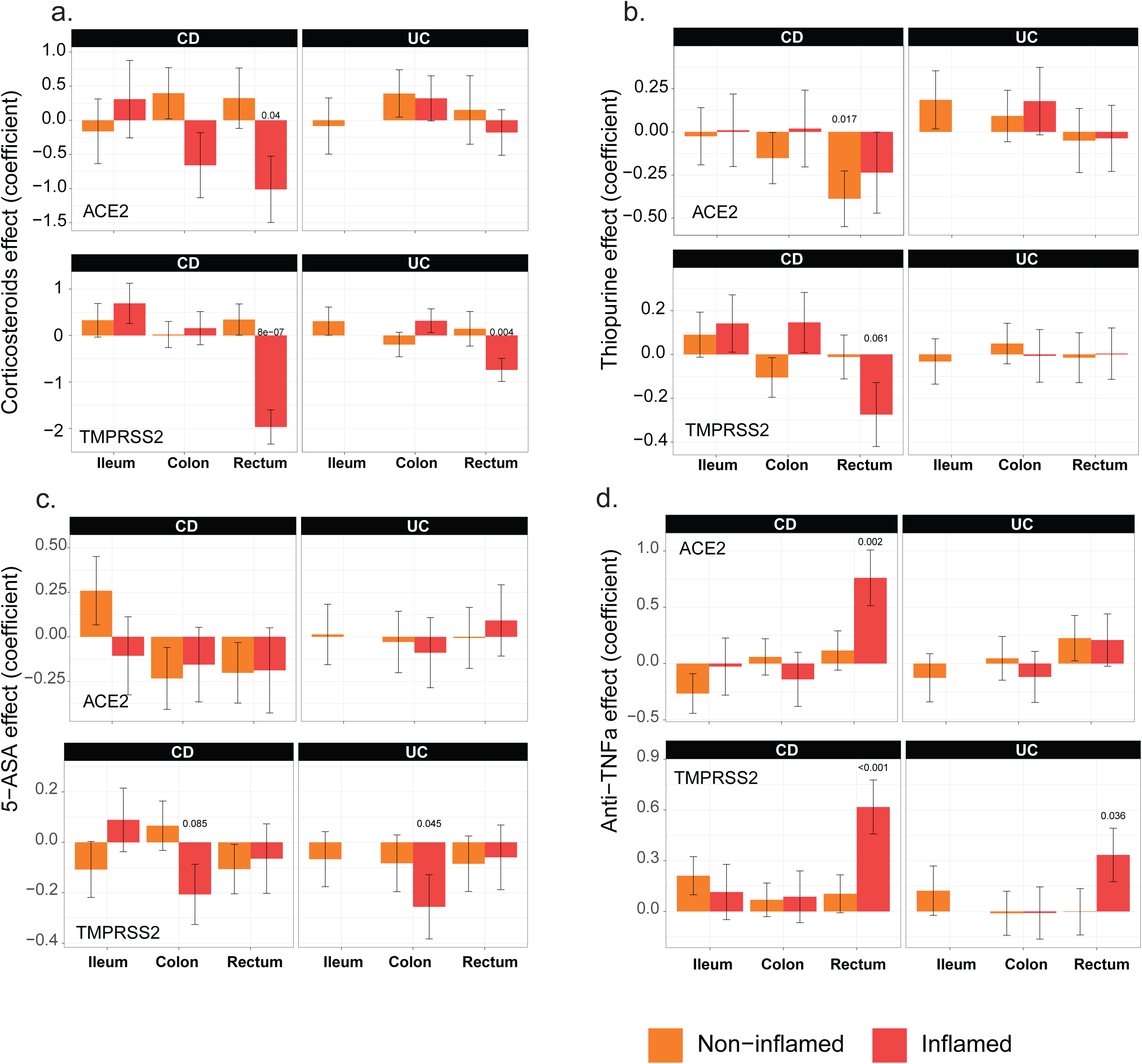
The effect of IBD medication use on expression of ACE2 and TMPRSS2 in intestinal biopsies of IBD and control MSCCR patients. Propensity-matched cohorts of UC and CD MSCCR patients receiving or not receiving corticosteroids (**A**), thiopurines (**B)**, 5-aminosalicylates (**C)**, or anti-TNF (**D)** were used to estimate the mean (+-SEM) differences in ACE2 (upper panel) and TMPRSS2 (lower panel) expression between the medicated and non-medicated groups. P-values <0.1 are reported. Under the model, we estimated the change in ACE2 or TMPRSS2 gene expression between the medicated and non-medicated group according to disease subtype (CD, UC) region (ileum, colon, rectum) and tissue type (inflamed, non-inflamed). Samples sizes are in Table S5.

### Among biologic medications, infliximab reduced ACE2 expression in the inflamed colon while ustekinumab increased ACE2 and TMPRSS2 in the inflamed colon in treatment responders

We also defined the effect of current anti-TNF therapy (either adalimumab or infliximab) use on ACE2 and TMPRSS2 (**Figure 3d**) expression. In the *ileum*, patients taking anti-TNF biologics did not show significantly different ACE2 or TMPRSS2 mRNA expression compared to those not on anti-TNF medication. In the large intestine, anti-TNF users showed increased ACE2 and TMPRSS expression, particularly in the inflamed rectum. Since the use cross-sectional cohort to study treatment effect has its limitations, we utilized published datasets from clinical trial cohorts, where longitudinal patient sampling was performed and information on endoscopic responses to treatment was available.

The results from a patient cohort treated with infliximab or vedolizumab on the GEMINI LTS trial^20^ are summarized (**Figure 4a and S5**). Compared to baseline ACE2 expression, post-infliximab (week 6) colonic ACE2 gene expression was significantly lower and this decrease was observed predominantly in endoscopic responders (**Figure S5**). In contrast, post-vedolizumab (week 6) ACE2 gene expression did not significantly change, although a nominally significant decrease was observed in endoscopic responders (**Figure 4a and S5**). Finally, neither vedolizumab nor infliximab modified TMPRSS2 expression.

**Figure 4:**
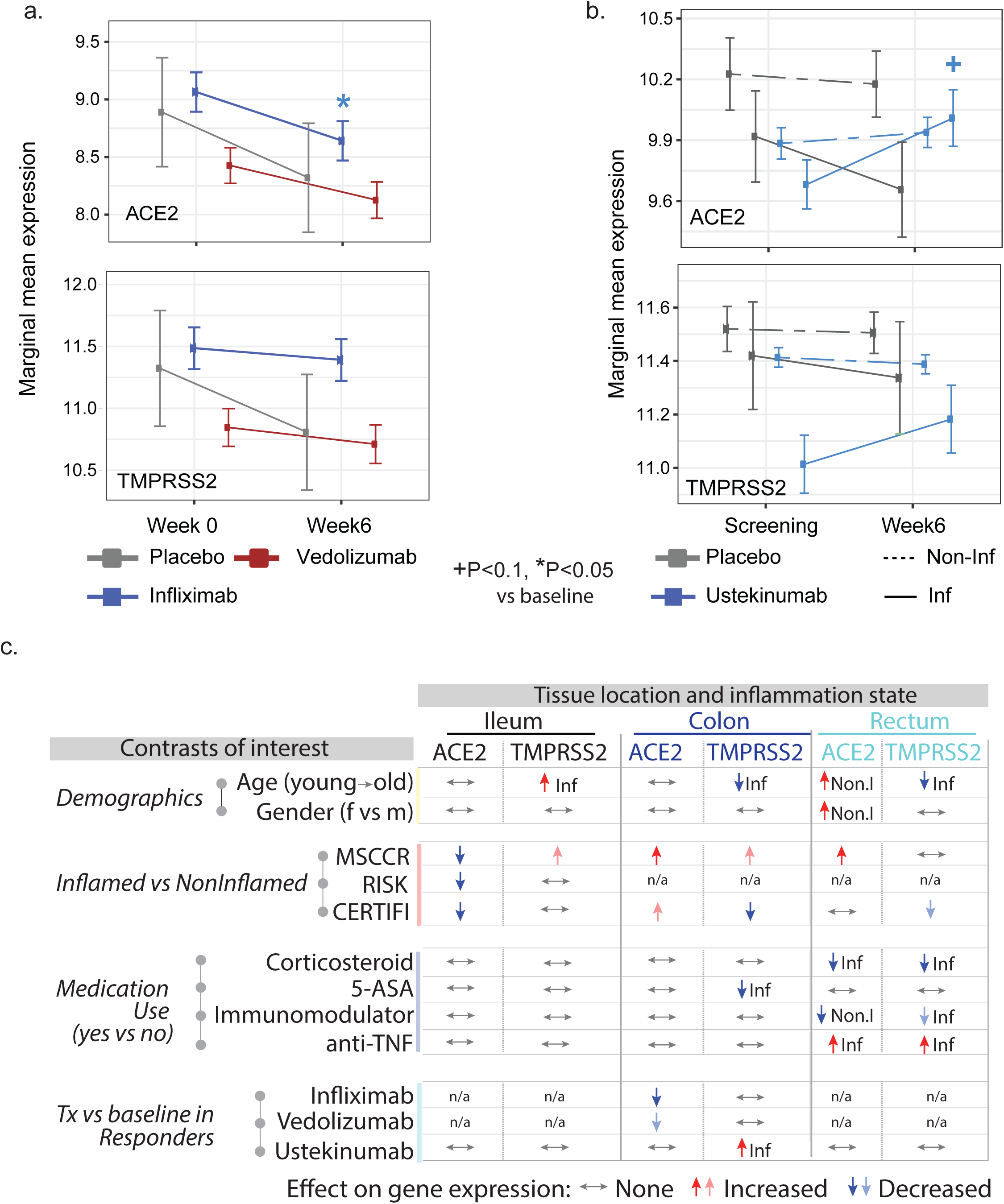
A summary of the effects of demographics, age, gender, inflammation and medication use on ACE2 and TMPRSS2 gene expression in the intestine. **A**) Changes on colonic gene expression profiles for ACE2 (upper panel) and TMPRSS2 (lower panel) on patients treated with vedolizumab (VDZ) or infliximab (IFX). **(A)** Estimated marginal means for the expression (M+-SEM) at baseline and week 4-6. **(B)** Changes in gut expression of ACE2 (upper panel) and TMPRSS2 (lower panel) in CD patients treated with ustekinumab (CERTIFI cohort). Estimated marginal means for the expression (M+-SEM) at baseline and week 6 for inflamed and non-inflamed biopsies. P-values denote significance of each time point compared to screening visit +(P<0.1), *(P<0.05), **(P<0.01). Samples sizes are in Table S6. **(C)** A summary of various contrasts of interest on ACE2 and TMPRSS2 expression in the ileum, colon or rectum. Arrows are used to indicate the effect of various conditions on ACE2 or TMPRSS2 gene expression, with Non.I indicating non-inflamed biopsy vs control and Inf indicating inflamed vs control. N/a indicates data not available. Lighter colored arrows represent nominal significance (p between 0.1 and 0.05) and darker arrows represent significance of p<0.05 in the various contrasts.

To study the impact of IL-12/IL-23-targeting ustekinumab on ACE2 and TMPRSS2 mRNA levels in IBD, we used the CERTIFI trial cohort^19, 27^. We first confirmed that, as in our MSCCR cohort, ACE2 gene expression was higher in uninflamed ileum compared to the colon regions (**Figure S6a**). Next, we observed that ileal ACE2 gene expression was significantly decreased with inflammation as was observed in the ileal samples from both MSCCR and RISK cohorts (**Figure S6a**). In colonic and rectal samples, a trend to increase ACE2 expression in inflamed biopsies as compared with non-inflamed was observed (**Figure S6a**).

Following ustekinumab treatment, ACE2 gene expression was increased (nominal significance) in the inflamed tissue (both small and large intestinal) after week 6 post ustekinumab as compared to the screening biopsy (**Figure 4b, Figure S6b**) contrasting with a decrease in expression in placebo-treated patients. Upon examining expression by ustekinumab response at week 22, the increase of ACE2 and TMPRSS2 expression was mainly observed in the colon and was stronger in responders (fold-change=1.79, p=0.055 for ACE2 and fold-change= 2.25, p=0.024 for TMPRSS2, **Figure S6c**). Thus, TNF-targeting biologics attenuated ACE2 mRNA expression in the inflamed colon, whereas IL-12/IL-23-targeting biologics increased both ACE2 and TMPRSS2 mRNA expression in the inflamed colon, particularly in therapy responders.

Overall, the effects of IBD medications on ACE2 and TMRSS2 mRNA is complex, region-specific and drug-specific. A summary of the medication effects is provided in Figure 4c.

### Intestinal ACE2 gene regulatory subnetworks are enriched in metabolic functions and interferon signaling in the inflamed colon

To identify potential functions associated with ACE2 and TMPRSS2 in the gut, we studied these genes in the context of Bayesian gene regulatory networks (BGRN). These probabilistic graphical models consider all available trait data (gene expression and genotype) simultaneously in order to derive gene:gene casual relationships amongst thousands of intermediate molecular traits^28^.

The nearest neighbors of ACE2 were extracted including genes within either 3 to 5 path lengths in each BGRN network (5 networks total) keeping the subnetwork sizes relatively similar (∼200-300 genes, **Figure 5a, 5d, Table S7**). Immediate neighbors of ACE2 in the ileum CD network included SLC6A19 (a known interacting partner of ACE2^29^), and other SLC transporters but also other viral-associated receptor proteins, like DPP4 and EZR^30^. A summary of the recurring genes across subnetworks is shown (**Figure 5b, 5e**).

**Figure 5:**
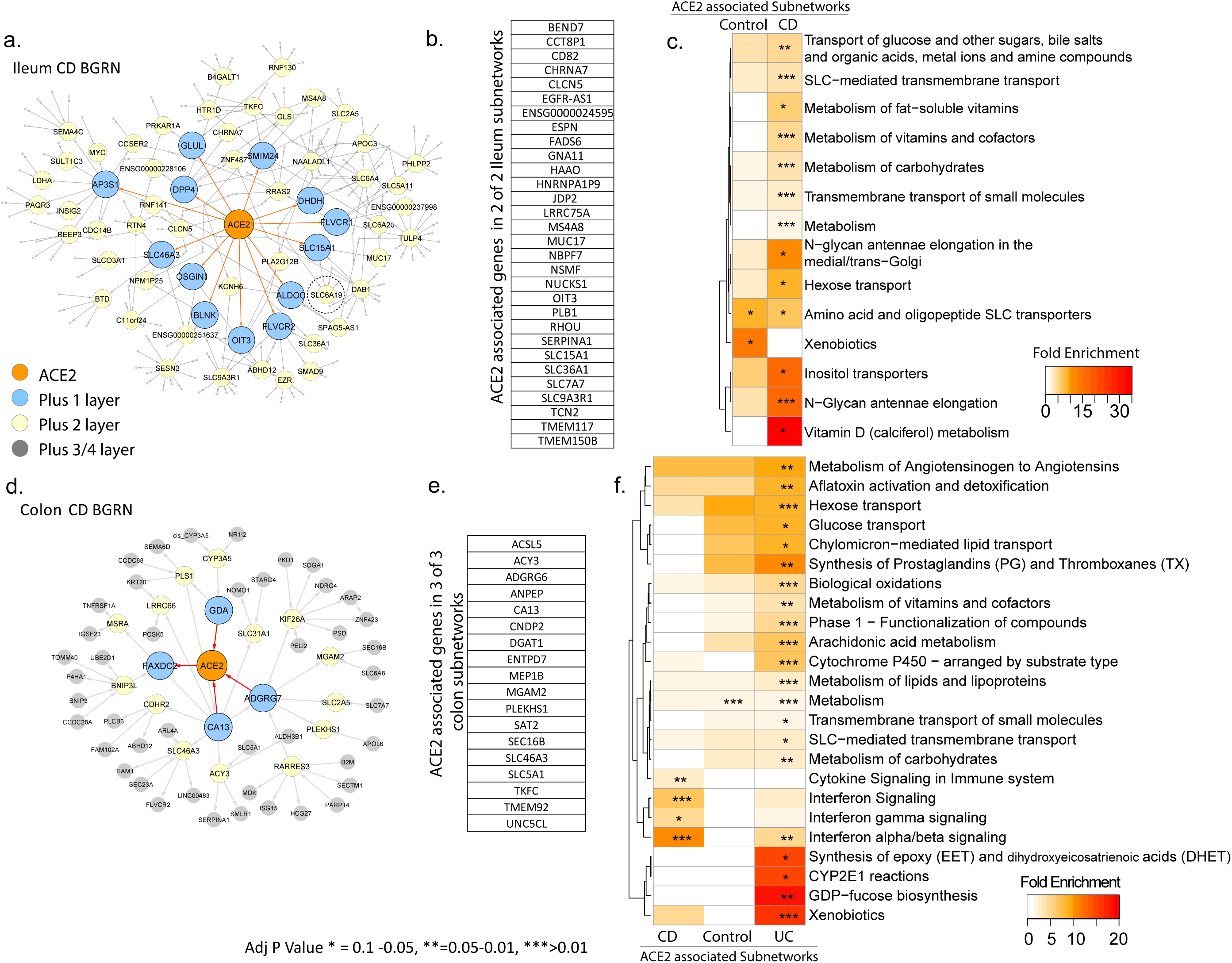
BGRN analysis of ACE2 reveals metabolic function and interferon pathway associations. The ACE2 associated subnetworks extracted from the **A**. Ileum CD BGRN (235 of 8458 nodes total) or from the (**D)** Colon CD BGRN (235 of 8549 total nodes). Genes (nodes) found within the first, second or third/fourth layer are colored blue, yellow or grey, respectively. Five ACE associated subnetworks were generated (Table S7). (**B)** and (**E)** A summary of the genes found in both ileum and colon ACE2-associated networks. Subnetwork sizes are: Control_ileum = 221; CD_ileum= 235; Control_colon = 229; CD_colon = 235 and UC_colon = 346. (**C)** and (**F)** Reactome pathway enrichment analysis of ACE2-associated subnetworks (with BH adj P <0.1). The heatmap depicts fold enrichment. Level of significance is indicated by either *, ** or *** for p-value of <0.1, <0.05 or <0.01, respectively (Table S8 for full results).

Functional enrichment of ACE2-associated subnetwork was interrogated in the ileum (**Figure 5c**) or colon (**Figure 5f**) using the Reactome database. Identified metabolic pathways included SLC-mediated transport; xenobiotic metabolism; vitamins and cofactors; and hexose transport. Additional ileum-associated pathways included amino acid and oligopeptide SLC transporters, while colon-associated networks included interferon and immune signaling.

To support our network approaches we verified a significant overlap was observed between the colonic ACE2-associated subnetworks and genes reported to be correlated with ACE2 expression in colonocytes^26^ (**Figure S7a**). In addition, ACE2-subnetworks were significantly enriched in expression profiles associated with epithelial cell types, including enterocytes and absorptive cells^23, 24^ (**Figure S7b-c**). Colonic ACE2-associated subnetworks also co-enriched in immune cell types as well as macrophage gene signatures following various cytokine perturbations including IFNγ/β^25^ (**Figure S7d**). In summary our ACE2-subnetworks are a novel source of insight into the regulation and function of ACE2.

### Intestinal TMPRSS2 gene regulatory subnetworks are predominantly enriched for metabolic functions

To address the function of TMRPSS2, the subnetworks associated with TMPRSS2 were extracted from the BGRNs (**Figure S8a**) allowing three or four layers to obtain similar subnetworks sizes (∼200–300 genes). TMPRSS2 was not found in the BGRN from the ileum of controls, likely due to a low expression variance, which is a filter used before BGRN network construction (**Table S13**). Genes recurring in 4 of 4 subnetworks were identified (**Figure S8b**). TMPRSS2-associated subnetwork genes are enriched for cell-cell communication, tight junction interaction, O-linked glycosylation of mucins and membrane trafficking-associated pathways (**Figure S8c**). Consistent with these functions, we observed enrichment of the TMPRSS2 subnetworks in genesets associated with enterocytes, goblet and secretory cells (**Figure S9 a-b**).

### A subset of pathways associated with SARS-CoV2 response and IBD inflammation overlap

COVID-19 is a multisystem disorder where innate and adaptive immune cells as well as non-immune cells likely play a role in disease pathogenesis. Therefore, apart from alterations in the receptor expression, we investigated additional areas of overlap between COVID-19 responsive pathways and pathways associated with IBD. As the first step, we examined a host molecular response signature generated following SARS-CoV-2 infection of a primary human lung epithelium (NHBE) cell and a transformed lung alveolar cell line (A549)^31^. Using GSVA, we generated a per-sample score summarizing expression of either up or down-regulated COVID-19-responsive genes, and evaluated differences in these scores according to intestinal region, disease and inflammation status (**Figure 6a and S10 a-b**). Genes up-regulated by SARS-CoV-2 infection show significantly higher expression in inflamed regions as compared to uninflamed regions or non-IBD control subjects, an observation confirmed in the CERTIFI CD cohort (**Figure S10c-d**). We directly compared the genes associated with response to lung cell SARS-CoV2 infection and various IBD-centric genesets generated in our MSCCR cohort^32^ or IBD GWAS genes. We observed that genes: up-regulated with inflammation, or positively associated with macroscopic or microscopic measures of disease, or associated with the risk of IBD, were significantly enriched with genes up-regulated by SARS-CoV2 infection of lung epithelial cells (**Figure S10e**).

**Figure 6:**
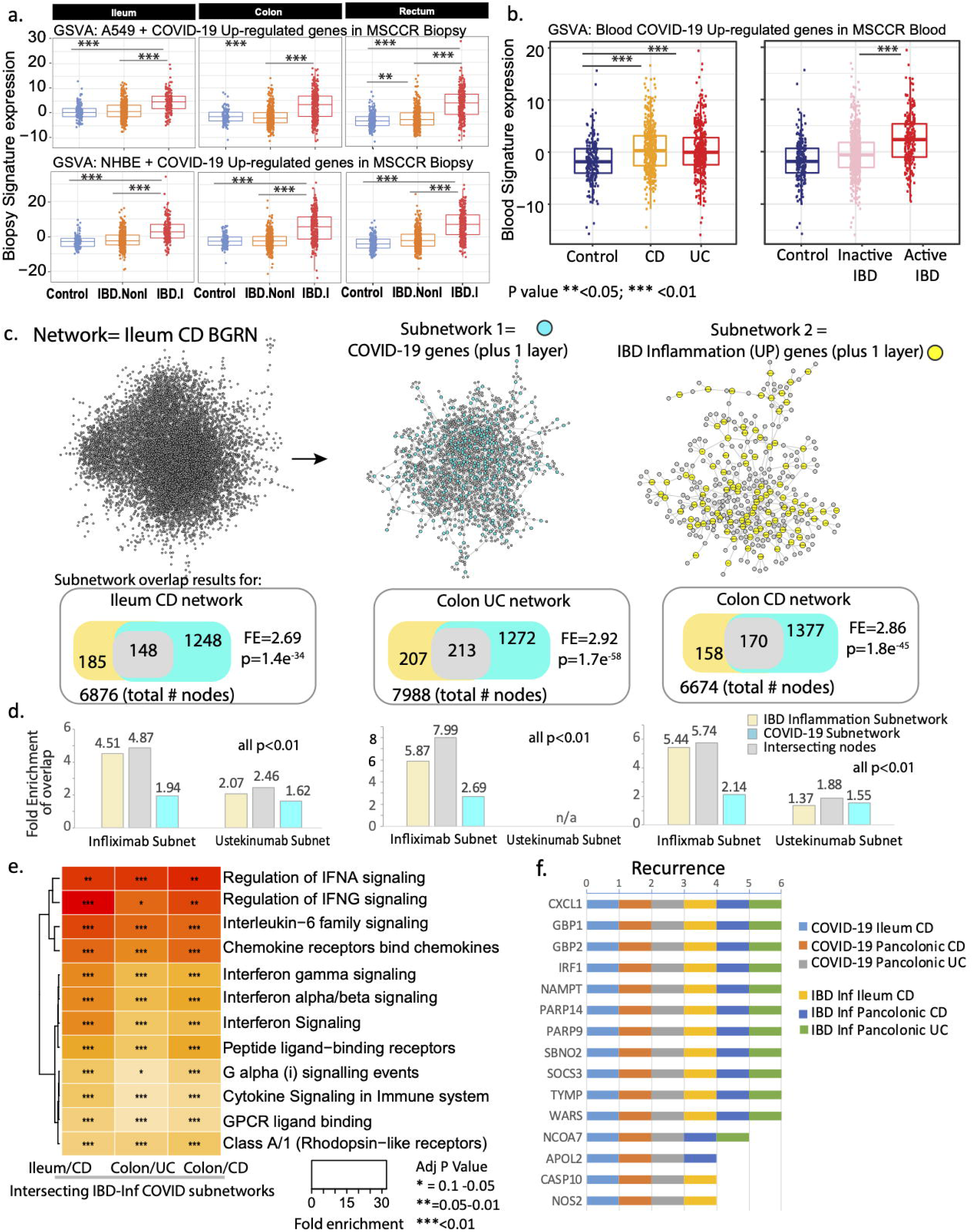
Molecular genes associated with IBD inflammation overlap with genes responsive to COVID-19 infection in lung cell model and whole blood. **(A)** Boxplot of the overall activity of COVID-19-responsive genes (up-regulated) as determined in A549 (upper panel) or NHBE (lower panel) lung epithelial cell models^31^ in the MSCCR biopsy samples using GSVA. Box plots summarize the mean expression (+/-SEM) of the two signatures (SF11 and sample sizes in Table S4, *** = p value <0.001). **(B)** Box plot summarizing the expression of the COVID-19 blood signature (up-regulated genes)^48^ in the blood transcriptome data of the MSCCR showing association between IBD disease (Left panel) and clinical severity (right panel) (SF11). **(C)** Three NHBE-COVID-19-and three IBD-Inf-associated subnetworks were generated one each for ileum CD, colon UC and colon CD BGRNs (Table S17 and S18). The three ileum-associated CD networks are shown for example. Venn diagrams of the overlap in subnetwork genes including the fold enrichment (FE) and a p-value of the enrichment test are shown. Using the BGRNs drug response subnetworks were generated (See supplementary methods) for infliximab and ustekinumab response genes. The bar graph summarizes the FE for the overlaps between the drug response and either the i) COVID-19 subnetworks; ii) IBD inflammation subnetworks or iii) the intersecting nodes between COVID-19 and IBD-Inf subnetworks. A full table of network nodes and enrichment results can be found in Table S23 and S24 **(D)** The 3 sets of genes in the intersecting subnetworks from C were interrogated for enrichment in the Reactome database. Heatmap depicting the FE in pathways (BH adj p<0.1 and minimum 5-fold enrichment, Table S19). **(E)** KDGs were determined, separately, for NHBE-COVID-19 or IBD-Inf signature genes in each BGRN and a summary of the top recurring KDGs are presented (Table S21 for full results). A gene was considered a KDG if its subnetwork (within 3 layers) was significantly enriched (Adj p<0.05) in signature genes.

Next, we examined for the congruence of COVID-19 related peripheral blood gene responses^32^ and active IBD inflammation. We observed that genes up-regulated in the blood of COVID-19 infected patients have significantly higher expression in the blood of IBD patients as compared to healthy control blood (**Figure 6b and Figure S11a**) as well as in patients with active IBD versus quiescent IBD (**Figure 6b and Figure S11b**).

To further interrogate the molecular congruence between SARS-CoV-2 infection and IBD inflammatory responses, we generated NHBE-COVID-19 associated subnetworks (NHBE-COVID_subnet) and IBD inflammation associated subnetworks (IBD_Inf_subnet) from three IBD BGRNs (**Table S17**). We then determined the overlap between them, with the rationale that common gene membership implies similar molecular pathobiologies. We observed a significant overlap between IBD-Inf and NHBE-COVID_subnets across all three independent networks tested (**Figure 6c**). Interestingly, the common genes (148 in ileum CD, 213 in Colon UC and 170 in Colon CD networks) were significantly enriched in candidate IBD GWAS genes (**Table S18**).

To determine the underlying shared patho-biological mechanisms we evaluated enrichment of the three *intersecting* subnetworks against Reactome pathways. Interestingly, although subnetworks were generated by projecting a COVID-19 response signature generated in an epithelial-based model the majority of the pathways and cell type enrichments were immune oriented with the most striking related to innate immune signaling via interferon and interleukin-6 (**Figure 6d, Figure S12, Table S19**). This is consistent with the fact that while SARS-CoV-2 infection initiates within the epithelium, COVID-19 is a multisystem disorder where innate and adaptive immune cells as well as non-immune cells likely play a role in disease pathogenesis. Our COVID-19 associated gut subnetwork findings are consistent with a large body of recent data that describe a dramatic up-regulation of pro-inflammatory cytokines in patients with COVID-19^33^ including induction of IFN-stimulated genes^31^.

We confirmed these observations using the recent dataset where SARS-CoV-2 was shown to infect hSIOs^34^ (**Figure S13a-b**). Direct overlap of genesets from the SARS-CoV-2 infected hSIOs and various IBD-centric genesets showed significant gene enrichments, similar to those observed between the lung-COVID-19 model and IBD (**Figure S13c**). Finally, subnetworks generated in the ileum CD using hSIO-COVID-19 responsive genesets (**Table S20**) overlapped with IBD-Inf associated subnetworks. The intersecting genes were similar to those found between IBD-Inf and NHBE-COVID19-associated ileum CD subnetworks, containing many interferon-stimulated genes (**Figure S13d-e**).

Next, as a data reduction approach, we evaluated each of the NHBE-COVID-19 and IBD-Inflammation molecular response subnetworks for key driver genes (KDGs). We then summarized the KDGs across the three BGRNs and determined those shared by IBD inflammation and NHBE-COVID-19 molecular responses (**Figure 6e, Table S21**). The shared KDGs identified were interferon-stimulated genes such as IRF1, GBP1/2, PARP14/9 and several of these were replicated in BGRNs from the RISK and CERTIFI cohort including CXCL1, GBP4, PARP9 amongst others (data not shown).

We also compared the intersections between COVID-19 associated genes and those associated with three different murine models of intestinal injury or inflammation. We observed a significant overlap between genes up-regulated with COVID-19 response in the NHBE lung model with genes upregulated in either a 1) Dextran Sodium Sulfate (DSS) induced intestinal injury model^35^; 2) TNBS-intestinal injury model^36^, especially two days post TNBS administration or 3) adoptive T cell transfer colitis model^37^ following a 6-week time period (ie W0<W2<W4<W6). Some examples of genes found commonly up-regulated amongst the 3 IBD-mouse models and NHBE-COVID-19 response included C3, IFITM3, IL1B, S100A9, TGM2 (transglutaminase 2) and PLAUR (plasminogen activator, urokinase receptor) (**Table S22**).

Finally, we generated colonic or ileal gene subnetworks associated with response to infliximab or ustekinumab (**Table S23**) and tested their enrichment in the tissue and IBD-type specific subnetworks and observed significant overlap between many of the genes associated with IBD therapy and either IBD-inflammation (used as positive control) or COVID-infection (**Figure 6c**) indicative of commonality between COVID-19 and response to IBD treatment. We also evaluated changes in the activity of the blood COVID-19 response signature in the blood of CERTIFI CD patients treated with ustekinumab and observed that COVID-19 up-regulated genes significantly decreased after 4 weeks of ustekinumab treatment (**Figure S14a**).

Altogether, through analyses of multiple genesets, we observed a significant overlap between COVID-19 response genes and genes associated with IBD response.

## Discussion

The objective of this study was to systematically determine the molecular intersections between COVID-19-associated inflammation, IBD and immunomodulatory drugs. Our data provides mRNA- and protein-level evidence of the regional distribution of ACE2 and TMPRSS2 across different parts of the GI tract and the impact of the commonly used IBD medications on ACE2 and TMPRSS2 expression in the inflamed and uninflamed intestines. Additionally, our data highlight an overlap between COVID-responsive pathways and pathways associated with IBD inflammation. These findings generate the possibility that some of the current and emerging therapies in IBD may be of benefit in patients with COVID-19.

In exploring the intersections between COVID-19 and IBD, we initially considered a) impact of inflammation on ACE2 expression and TMPRSS2 expression in the intestines; b) impact of IBD medications on ACE2 and TMPRSS2 expression. ACE2 has a less appreciated RAAS-independent role in the intestine by promoting amino acid absorption. Consistent with this function we observed high levels of ACE2 expression in the small bowel brush border^38-40^ supportive of the role of ACE2 in mucosal homeostasis^39^. Remarkably, the analyses of gene:regulatory networks empirically derived from the terminal ileum, showed that ACE2 was co-regulated with SLC6A19, an amino acid transporter that physically interacts with ACE2^41^. The colon-derived ACE2 subnetwork would also suggest, a potential metabolic role for ACE2 in the colon including solute carrier dependent processes.

While the physiological function of TMPRSS2 remains largely elusive, it has been linked to epithelial sodium channel (ENaC) regulation^42^ and possibly in regulating sperm function^43^. In agreement with literature^42^, our BGRN analysis showed SCNN1A, which corresponds to a subunit of the ENaC, associated with TMPRSS2. Remarkably, gene products co-localized with TMPRSS2 were consistently enriched in fundamental epithelial functions. For example, F11R, which encodes a JAM-A protein mediates tight junction formation. Interestingly, we noted connections to viral responses. JAM-A is a reovirus receptor^44^ and TRIM31^45^ and BAIAIP2L1^46^ associate with mitochondrial anti-viral associated proteins.

We have observed a reduction in ACE2 expression in the inflamed ileum. Since ACE2 appears to be a brush-border enzyme, its reduction with inflammation in the ileum is consistent with loss of expression of other brush border enzymes during enteritis^47^. That said, even during inflammation, the expression of ACE2 remains significant in the small intestines. In contrast, inflammation associated with IBD enhances the expression of ACE2 and TMPRSS2 in the rectum. Overall, modulation of ACE2 and TMPRSS2 by IBD-associated inflammation is complex and appears to be region-specific in the intestines.

Akin to the impact of inflammation on intestinal ACE2 and TMPRSS2 expression, the effect of IBD medications was complex. Corticosteroids, thiopurines or 5-aminosalicylates did not significantly affect TMPRSS2 mRNA expression in the ileum. However, each of these three medications significantly decreased TMPRSS2 mRNA expression in inflamed rectum or colon samples.

Importantly, while SARS-CoV-2 infection initiates with the viral attachment to ACE2 and its cleavage by TMPRSS2 within the epithelial surfaces^7-10^, COVID-19 is a multisystem disorder where innate and adaptive immune cells as well as non-immune cells likely play a role in disease pathogenesis. Therefore, apart from alterations in the receptor expression, we investigated additional areas of overlap between COVID-19 responsive pathways and pathways associated with IBD.

Notably, a number of such intersections appeared. For example, *IL6, CXCL1/2/5, PDPN, S100A8/A9* which were upregulated following SARS-CoV-2 infection of primary human lung epithelium (NHBE)^31^ were also significantly upregulated in the inflamed intestines of patients in our IBD cohort (MSCCR). Similarly, a peripheral blood gene signature from COVID-19 patients^48^ which included the upregulated genes, *CLEC4D, S100A8/A9* and *FCAR* (aka CD89), were also significantly upregulated in the blood of our MSCCR patients with IBD (as compared to controls) and in patients with active IBD (as compared to patients with quiescent disease). Additionally, a number of SARS-CoV-2-associated genes were upregulated in murine models of intestinal injury with DSS^35^ (*DAPP1, PDPN, IL1RN, DUOX2, IL1B, S100A9, CXCL2, CXCL3*) or TNBS^36^ (*MARCKSL1, IFITIM3, IFITIM1, C3, AGR2, REG4*) or adoptive T cell transfer colitis model^37^ (*TAP2, MARCKSL1, SLP1, PARP9, IFITIM3, MMP13, IL1B, S100A8*). These genes relate to a number of IBD-relevant pathways including those associated with inflammatory cytokine signaling (including IL-6, IL-1, IFN-G), chemokine signaling, but also with interferon-associated pathways, regulation of complement cascade, G protein coupled receptor (GPCR) signaling as well as collagen degradation.

Having observed significant molecular intersections between COVID-19- and IBD-associated pathways, we next examined the impact of biologic medications used in IBD therapeutics where pre- and post-treatment transcriptomic data was available. Interestingly, our network analyses had identified a number of shared COVID-19 and IBD-associated ‘key driver genes’, including *CXCL1, GBP4, SOCS3, PARP14* and *PARP9*. Importantly, we observed that these KDGs which were up-regulated with COVID-19 and IBD inflammation, were all down-regulated following infliximab treatment. Thus, the impact of IBD medications on attenuating some of the key inflammatory genes and pathways would be independent of their complex effects on the expression of ACE2 and TMPRSS2 on enterocytes.

Given the unregulated inflammatory responses in patients with severe COVID-19, it has been argued that targeted use of anti-inflammatory medications be considered as a therapeutic option^49^. This approach has been met with variable success. While the use of dexamethasone has provided a striking mortality benefit^50^, the results of trials using anti-IL-6 have been equivocal (NCT04320615) and NCT04372186. An anti-TNF trial is currently underway in the UK (NCT04425538). Our data suggest that anti-IL-12/23 therapy could also be considered in the therapeutic armamentarium in patients with COVID-19. Reassuringly, real-time data from a registry of IBD-COVID patients^51^ did not report significant adverse outcomes associated with the use of biologic medications including anti-TNF and anti-IL-12/23 inhibitors. To the contrary, the risk of severe COVID-19 was found to be reduced in IBD patients on anti-TNF inhibitor medication, although the results of such population-based studies should be interpreted with caution^51^.

In summary, through detailed analyses of intestinal tissues in health and IBD, we conclude that high expression of ACE2 and TMPRSS2 potentially supports local, GI-associated replication of SARS-COV-2. Further, a number of overlapping inflammatory pathways between COVID-19 and IBD are noted. These data support the use of specific anti-inflammatory agents in the treatment of patients with COVID-19.

**Author names in bold designate equal contribution**

**Author names in bold designate equal contribution**

## ACKNOWLEDGMENTS

We thank the patients who participated in the study.

## Funding

This work was supported by the following grants: NIH/NIDDK R01 112296 (SM). Additional support was provided by K23KD111995 (RCU) and a Career Development Award from the Crohn’s and Colitis Foundation (RCU). CA, MSF were supported by a Litwin Pioneers award grant. CA and ES were supported in part by the Helmsley Charitable Trust. MT was supported by the Digestive Disease Research Foundation (DDRF). The sampling of the Inflammatory Bowel Disease cohort (Crohn’s disease and ulcerative colitis) was jointly designed as part of the research alliance between Janssen Biotech, Inc. and The Icahn School of Medicine at Mount Sinai. Beyond this exception, no other funders had a role in analyses design and interpretation. This work was supported in part through the computational resources and staff expertise provided by Scientific Computing at the Icahn School of Medicine at Mount Sinai.

## Competing interests

SM and JFC have an unrestricted, investigator-initiated grant from Takeda Pharmaceuticals to examine novel homing mechanisms to the GI tract. RCU has served as an advisory board member or consultant for Eli Lilly, Janssen, Pfizer and Takeda. Mount Sinai co-authors (from Genetics and Genomics, Icahn Institute for Data Science and Genomic Technology, Human Immune Monitoring Center, Population Health Science and Policy, Division of Gastroenterology, Pediatric GI and Hepatology, Susan and Leonard Feinstein IBD Clinical Center at Icahn School of Medicine at Mount Sinai) were partially funded as part of research alliance between Janssen Biotech and The Icahn School of Medicine at Mount Sinai. MC, AS, JP and CB are employees at Research and Development. and JRF is a former employee at Janssen Research and Development and is currently employed at Alnylam Pharmaceuticals. MD is a consultant for Janssen. BES discloses consulting fees from 4D Pharma, Abbvie, Allergan, Amgen, Arena Pharmaceuticals, AstraZeneca, BoehringerIngelheim, Boston Pharmaceuticals, Capella Biosciences, Celgene, Celltrion Healthcare, EnGene, Ferring, Genentech, Gilead, Hoffmann-La Roche, Immunic, Ironwood Pharmaceuticals, Janssen, Lilly, Lyndra, MedImmune, Morphic Therapeutic, Oppilan Pharma, OSE Immunotherapeutics, Otsuka, Palatin Technologies, Pfizer, Progenity, Prometheus Laboratories, Redhill Biopharma, Rheos Medicines, Seres Therapeutics, Shire, Synergy Pharmaceuticals, Takeda, Target PharmaSolutions, Theravance Biopharma R&D, TiGenix, Vivelix Pharmaceuticals; honoraria for speaking in CME programs from Takeda, Janssen, Lilly, Gilead, Pfizer, Genetech; research funding from Celgene, Pfizer, Takeda, Theravance Biopharma R&D, Janssen. MCD discloses consulting fees from Abbvie, Allergan, Amgen, Arena Pharmaceuticals, AstraZeneca, BoehringerIngelheim, Celgene, Ferring, Genentech, Gilead, Hoffmann-La Roche, Janssen, Pfizer, Prometheus Biosciences, Takeda, Target PharmaSolutions and research funding from Abbvie, Janssen, Pfizer, Prometheus Biosciences Takeda. Dr. Colombel reports receiving research grants from AbbVie, Janssen Pharmaceuticals and Takeda; receiving payment for lectures from AbbVie, Amgen, Allergan, Inc. Ferring Pharmaceuticals, Shire, and Takeda; receiving consulting fees from AbbVie, Amgen, Arena Pharmaceuticals, Boehringer Ingelheim, Celgene Corporation, Celltrion, Eli Lilly, Enterome, Ferring Pharmaceuticals, Geneva, Genentech, Janssen Pharmaceuticals, Landos, Ipsen, Imedex, Medimmune, Merck, Novartis, O Mass, Otsuka, Pfizer, Shire, Takeda, Tigenix, Viela bio; and holds stock options in Intestinal Biotech Development and Genfit.

## Supplementary Legends

**Figure S1. TMPRSS2 and ACE2 distribution and localization in the intestine in children and adults**. Representative immunofluorescence images of ACE2 (green) and TMPRSS2 (red) counterstained with DAPI (blue) in intestinal biopsies of non-IBD patients. Magnified images of surface epithelium (se) and crypt epithelium (ce) showing only TMPRSS2 and DAPI. (**A)** Duodenal biopsies (**B)** Terminal ileum biopsies (**C)** Biopsies from indicated colonic segments. Patient age (years) and sex (M, male; F, female) indicated in the top left corner of each image. Isotype controls and no primary controls for each segment are included on the far right of each panel. Scale bar, 100µm.

**Figure S2: Expression of ACE2 and TMPRSS2 protein in the intestine of IBD patients. (A)** Representative immunofluorescence images of ACE2 (green) and DAPI (blue) on the left and TMPRSS2 (red) and DAPI (blue) on the right in paired inflamed and uninflamed IBD intestinal specimens. Terminal ileum from a CD patient pre-IFX (inflamed) and after IFX treatment (uninflamed). Inflamed left colon and uninflamed sigmoid colon from a UC patient pre-biologic. Inflamed rectum from a CD patient pre-IFX and uninflamed rectum post-IFX therapy. Clinical characteristics of IBD patients and biopsies are summarized in supplementary table 3.

**Figure S3:** ACE2 and TMPRSS2 gut gene expression in MSCCR CD and UC patients versus healthy controls. Plots summarize the expression level of ACE2 (**A**) and TMPRSS2 (**B**) across various gut regions from CD (top panel) or UC (bottom panel) which were either endoscopically non-inflamed (Non-Inf) or inflamed (Inf). Each IBD region is compared to healthy controls. Numbers at the bottom represent the number of samples (#Non-Inf vs #Control / #Inf vs #Control). Level of significance is indicated by either *, ** or *** for p-value <0.05, <0.01 or <0.005, respectively.

**Figure S4: ACE2 and TMPRSS2 gene expression according to location of disease in CD patients and effects of age and gender. (A)** Normalized gene expression of ACE2 (top panel) and TMPRSS2 (bottom panel) summarized according to CD patients with L1 (purple) or L2/L3 (orange) disease separated by region. Biopsies from endoscopically defined non-inflamed (left panel) or inflamed (right panel) areas are examined separately. (**B)** The effect of gender and age effect on ACE2 TMPRSS2 gene expression was estimated using a multivariable model with smoking, age, gender and two interaction of age and gender and IBD subtype and Tissue. The coefficient for Age effect is presented for CD and UC at different regions (ileum, colon and rectum). P values denote significance of the age effect for each disease and tissue group, +(P<0.1), *(P<0.05), **(P<0.01). (**C)** We estimated the marginal mean expression of ACE2 and TMPRSS2 for male and female at each region (ileum, colon and rectum) and disease (CD and UC) group. P values are presented where significant differences between males and females were found.

**Figure S5: The effect of infliximab and vedolizumab on ACE2 and TMPRSS2 expression. A**) Changes on colonic gene expression profiles for ACE2 (upper panel) and TMPRSS2 (lower panel) on patients treated with vedolizumab (VDZ) or infliximab (IFX). Differences in endoscopic responders and non-responders at week 4-6 versus baseline samples. (P-values denote significance of each time point compared to screening visit +(P<0.1), *(P<0.05), **(P<0.01). Samples sizes are in Table S6.

**Figure S6: The effect of ustekinumab (CERTIFI cohort) on ACE2 and TMPRSS2 gene expression in the intestine**. (**A**) Baseline differences in expression of ACE2 and TMPRSS2 between Inflamed and Non-inflamed tissue was estimated across different regions (Ileum, Colon and Rectum) using a mixed-effect model with Tissue and Region and its interactions as fixed effects and random intercepts for each patient. P values indicated the significance of the Inflamed vs. Non-Inflamed comparison. (**B)** Changes in gut expression of ACE2 (upper panel) and TMPRSS2 (lower panel) in CD patients treated with ustekinumab (CERTIFI cohort). Treatment changes in expression of ACE2 and TMPRSS2 were modeled using a mixed-effect model with visit, region, tissue and treatment and its interactions as fixed effect. Marginal estimated means are presented for patients treated with ustekinumab and placebo group at baseline and week 6 across different gut regions. P values denote significance of change at Week 6 from screening, +(P<0.1), *(P<0.05). **C)** As no change was observed in non-inflamed biopsies, treatment effect on inflamed biopsies was compared between week-22 clinical responders and non-responders versus baseline.+(P<0.1), *(P<0.05). The samples sizes are summarized in Table S6

**Figure S7: ACE2 associated subnetworks reveals gut epithelial cell type enrichment**. Each ACE2 associated subnetwork was interrogated for enrichment with: (**A)** Genes co-correlated with ACE2 in colonocytes was curated from Wang et al^26^; (**B)** Gut cell type associated signatures from (Huang et al.,2019^24^) and (**C)** (Smillie et al., 2019^23^). (**D)** Gene sets associated with various perturbations in macrophages^25^. The number of genes within each ACE2 associated subnetwork are: Control ileum = 221; CD ileum= 235; Control colon = 229; CD colon = 235 and UC colon = 346. The heatmap coloring depicts the fold enrichment and the level of significance based on BH adj p-value of either *(P<0.1), **(P<0.05) or ***(P<0.01). The full enrichment results are available in Tables S9 to S12.

**Figure S8: Bayesian gene regulatory network (BGRN) analysis of TMPRSS2 reveals gut barrier function pathway associations. (A)** The TMPRSS2 subnetwork extracted from the UC Colon BGRN (490 of 8557 nodes total) was generated by including 4 additional layers of genes (undirected network expansion). Genes or nodes found within the first, second or third/fourth layer are colored blue, yellow or grey, respectively. In total 4 TMPRSS2 subnetworks were generated, one from ileum gene expression data from MSCCR CD patients and 3 from colon (and rectum) gene expression data from MSCCR control, UC, or CD patients (See Table S13). Both inflamed and non-inflamed biopsies used for the IBD networks are shown. (**B)** A summary of the genes found in 4 out of the 4 TMPRSS2 associated networks. The number of genes within each TMPRSS2 associated subnetworks are: CD ileum= 397; Control colon= 268; CD colon = 366 and UC colon= 490. (**C**) Pathway enrichment analysis (Fisher’s exact test) of each TMPRSS2 associated subnetwork according to Reactome pathways was performed. Only pathways which were found significantly enriched in at least one TMPRSS2 associated subnetwork (at BH adj P <0.1) are presented in the heatmaps in C. The heatmap coloring depicts the fold enrichment and the level of significance based on BH adj p-value of either *(P<0.1), **(P<0.05) or ***(P<0.01). The full enrichment results are available in Table S14.

**Figure S9: TMPRSS2 associated subnetworks reveals gut cell type enrichment**. Each of the four TMPRSS2 associated subnetworks were examined for enrichment with various gut cell type related signatures^23, 24^ (Smillie et al.,2019) (**A**) (Huang et al., 2019) (**B**) The number of genes within each TMPRSS2 associated subnetworks are: CD ileum= 397; Control colon= 268; CD colon = 366 and UC colon= 490. The heatmap coloring depicts the fold enrichment and the level of significance based on BH adj p-value of either *(P<0.1), **(P<0.05) or ***(P<0.01). The full enrichment results are available in Tables S15 and S16.

**Figure S10: Geneset variation analysis of lung COVID-19 responsive genes as determined in MSCCR and CERTIFI cohort gut biopsies. A**. We evaluated expression of COVID-19-responsive genes as determined in NHBE (**A**) or A549 (**B**) lung epithelial cell models^31^ in the MSCCR biopsy samples using GSVA. Mixed-effect linear models with Region, tissue type and its interaction were used to compare COVID-19 scores between control, non-inflamed and inflamed samples (Sample sizes in Table S4, *** indicates p value <0.001). (**C, D**) Two molecular expression signatures reflecting a host’s transcriptional response to SARS-CoV-2 infection were curated from Blanco-Melo et al^31^. (**C)** Transformed lung alveolar cell line (A549) and (**D)** Primary human lung epithelium (NHBE) were profiled following SARS-CoV-2 exposure. Boxplots for the GSVA scores of COVID-19-responsive genes in inflamed and non-inflamed biopsies across gut regions for patients in CERTIFI cohort. Mixed-effect models with region and tissue as fixed-effects were used to compare expression between non-inflamed and inflamed samples. Sample sizes are presented in Table S6. (**E)** A heatmap summarizing the significance (-log adj P value) for the enrichment of genes Up-or Down-regulated following lung cell SARS-CoV2 infection in various IBD disease associated genesets derived from the MSCCR cohort analysis and IBD GWAS genes.

**Figure S11: Expression of a blood signature of genes responsive to COVID-19 infection in blood transcriptome of adult IBD patients and controls patients (MSCCR cohort)**. Box plot summarizing the expression of the blood signature identified in COVID-19 infected patients versus healthy controls^48^ as determined in the blood transcriptome data of A. the MSCCR CD (n=432) and UC (n=389) patients and controls (n=209) and B. between clinically-defined inactive (n=288 CD, n=340 UC) versus active (n=72 CD, n=49 UC) MSCCR IBD patients (right panel). GSVA scores associated with up-regulated genes are shown in the left panel and down-regulated genes are shown in the right panel. P values denote significance with *P<0.05, **P<0.01 and ***P<0.001.

**Figure S12: Lung model COVID-19 associated gene subnetworks and the shared subnetwork genes with IBD-inflamed genes are enriched with immune cell types**. NHBE COVID-19 subnetworks and the intersecting genes found between them and the IBD-inflamed subnetwork genes were interrogated for enrichment with gut cell type signatures (Smillie et al., 2019)^23^. The heatmap coloring depicts the fold enrichment and the level of significance based on BH adj p-value of either *(P<0.1), **(P<0.05) or ***(P<0.01).

**Figure S13: Subnetworks associated with COVID-19 response in hSIOs overlaps with NHBE-COVID-19 responsive subnetworks**. Two molecular expression signatures reflecting transcriptional responses to SARS-CoV-2 infection in human small intestinal organoids (hSIOs) were curated^34^. **(A)** hSIOs cultured with differentiation media (DIF) contain predominantly enterocytes, goblet cells and low numbers of enteroendocrine cells. **(B)** hSIOs grown in Wnt high expansion medium (EXP) consist mainly of stem cells and enterocyte progenitors. We determined expression of the hSIO COVID-19-responsive genes (at FDR<0.1) in the ileum, colon and rectum MSCCR cohort samples using geneset variation analysis (GSVA) and mixed-effect model with region, tissue type and its interaction as fixed effects was performed to compare expression between control, non-inflamed and inflamed samples. P values are as indicated. **(C)** A heatmap summarizing the significance (-log adj P value) for the enrichment of genes Up- or Down-regulated following hSIO SARS-CoV2 infection in various IBD disease associated genesets derived from the MSCCR cohort analysis and IBD GWAS genes. **(D)** A Venn diagram summarizing the overlaps of 3 subnetworks generated using the ileum CD BGRN, from projecting signatures associated with either 1. IBD-Inf; 2. NHBE_COVID-19 response or 3: hSIO (DIF media)-COVID-19 response, allowing 1 or 2 nearest neighboring genes. **(E)** A Venn diagram summarizing the overlaps of 3 subnetworks generated using the ileum CD BGRN, from projecting signatures associated with either 1. IBD-Inf; 2. NHBE_COVID-19 response or 3: hSIO (EXP media)-COVID-19 response, allowing 1 nearest neighboring gene. The hSIO(DIF)- and hSIO(EXP)-COVID-19 associated subnetworks are found in Table S20. The intersecting genes are shown. Note many genes are also identified as KDGs in Figure 6e. The significance of the overlaps of various genesets is presented in each panel.

**Figure S14: Expression of a molecular signature consisting of genes responsive to COVID-19 infection as determined in blood transcriptome data from adult IBD patients with CD (CERTIFI cohort). (A)** Estimated marginal means (M+SEM) for the activity (GSVA scores) of the COVID-19 blood signature (COVID-19 infected patients versus healthy controls)^48^ on the blood transcriptome of CERTIFI patients with CD at baseline and after weeks 4 to 22 of treatment with ustekinumab or placebo. (**B)** Ustekinumab-induced changes in COVID-19 blood signature between responders and non-responders (defined as change in CDAI). P-values denote significance of each time point compared to screening visit *P<0.1), **(P<0.05), ***(P<0.01).

## Supplementary Table Legends

**Table S1:** Clinical characteristics of non-IBD patients included in immunofluorescence analysis in Figure 1. Abbreviations: TI, terminal ileum; CRC, colorectal cancer; DM2, type 2 diabetes mellitus; HLD, hyperlipidemia; HTN, hypertension; GERD, gastroesophageal reflux disease; ED, erectile dysfunction; PUD, peptic ulcer disease; OA, osteoarthritis; BPH, benign prostatic hyperplasia; CAD, coronary artery disease; IBS, irritable bowel syndrome.

**Table S2**: Clinical characteristics of IBD patients included in immunofluorescence analysis in Figure S1. Clinical characteristics of IBD patients before (a) and/or after (b) biologic therapy. Macroscopic extent of disease refers to the Montreal classification for UC patients. Biologic naive is defined as no previous exposure to biologics. Abbreviations: SES-CD, Simple endoscopic score for Crohn Disease; 6MP, mercaptopurine; HBV, hepatitis B; HTN, hypertension; Afib, atrial fibrillation; CAD, coronary artery disease; GERD, gastroesophageal reflux disease; PPROM, preterm premature rupture of membranes; TI, terminal ileum.

**Table S3:** Clinical characteristics of IBD patients from the Mount Sinai Crohn’s and Colitis Registry (MSCCR).

**Table S4:** Sample sizes associated with analysis in Figure 2a.

**Table S5:** Sample sizes associated with analysis in Figure 3.

**Table S6:** Sample sizes associated with analysis in Figure 4 and Figure S3.

**Table S7**: Subnetworks associated data in Figure 5a and 5d.

**Table S8:** Data associated with Figure 5c and f

**Table S9:** Data associated with Figure S5a

**Table S10:** Data associated with Figure S5b

**Table S11:** Data associated with Figure S5c

**Table S12:** Data associated with Figure S5d

**Table S13:** Data associated with Figure S6a

**Table S14:** Data associated with Figure S6c

**Table S15:** Data associated with Figure S7a

**Table S16:** Data associated with Figure S7b

**Table S17:** Data associated with Figure 6c networks

**Table S18:** Data associated with Figure 6c

**Table S19**: Data for Figure 6d

**Table S20:** Data for Figure S9d and e

**Table S21:** Data for Figure 6e (KDGs)

**Table S22:** Results of enrichment test between COVID-19 response genes (in NHBE model) and Mouse IBD model differentially expressed genes

**Table S23:** Drug response associated CD ileum subnetworks

