## Supplementary Methods for "Intestinal inflammation modulates the expression of ACE2 and TMPRSS2 and potentially overlaps with the pathogenesis of SARS-CoV-2 related disease"

**Immunofluorescence microscopy:**

Specimens were obtained via clinical endoscopy during routine care (**Table S1 and S2**). Tissue was formalin fixed and paraffin embedded by the clinical pathology core at our institution. Bowel sections (5µm) were dewaxed in xylene and rehydrated in graded alcohol followed by 2 washes in 0.01M phosphate-buffered saline (PBS). Antigen-retrieval was performed by heating slides in a pressure cooker in target retrieval solution (Dako, S1699) for 15 minutes. After washing 3 times in PBS, non-specific binding was blocked by 5% normal donkey serum in PBST (0.1% tween 20, PBS) for 45 min at room temperature. Sections were then incubated in primary antibodies diluted in blocking solution overnight at 4°C. Primary antibodies used included ACE2 (abcam, ab15348, 1:1000) and EPCAM (abcam, ab228023, prediluted).

Sections were washed 3 times in PBS and incubated in secondary antibody (Alexa Flour 488 donkey anti-rabbit, Alexa Flour 594 donkey anti-mouse) and 4',6-diamidino-2-phenylindole (DAPI) diluted in PBS for 1 hour at room temperature. Sections were washed three times in PBS and mounted with fluoromount G (Electron microscopy sciences, 1798425). Controls included, omitting primary antibody (no primary control), or substituting primary antibodies with non-reactive polyclonal rabbit IgG (abcam, ab37415) and/or monoclonal mouse IgG1 kappa (abcam, ab170190) antibodies (isotype control).  Slides were visualized and imaged using a Nikon Eclipse Ni microscope and digital SLR camera (Nikon, DS-Qi2).

Due to the low expression of TMPRSS2 protein in the gut, Tyramide SuperBoost Kit (Thermofisher, B40915) was used to amplify the signal. After dewaxing, rehydration and antigen retrieval (as above), endogenous HRP was quenched by incubating slides in 3% hydrogen peroxide for 1 hour at room temperature. The slides were then washed in PBS 3 times and blocked in 10% goat serum for 1 hour at room temperature. Slides were then incubated in mouse anti-TMPRSS2 (Millipore, MABF2158, 1:500) diluted in blocking buffer at 4°C overnight. After rinsing 3 times in PBS, sections were incubated in poly-HRP-conjugated goat anti-mouse secondary antibody for 1 hour at room temperature.  Slides were incubated in tyramide working solution for 10 minutes before the reaction was stopped. Sections were rinsed with PBS 3 times and then probed for other antigens as described above except goat serum and goat secondary antibodies were used instead of donkey.

**Analysis of Mount Sinai Crohn’s and Colitis Registry (MSCCR) Cohort:**

Biopsy RNA was extracted and processed in randomly allocated batchers as previously described^1^. RNA was isolated from frozen tissue using Qiagen QIAsymphony RNA Kit (cat.# 931636) on the QIAsymphony.  RNA from whole blood collected in PAXgene tubes was isolated using QIAsymphony Blood PAXgene RNA kit (cat.# 762635).  One microgram of total RNA was used for the preparation of the sequencing libraries using the RNA Tru Seq Kit (Illumina (Cat # RS-122-2001-48). Ribosomal RNA from biopsy tissue was depleted from total RNA using the Ribozero kit (Illumina Cat # MRZG12324), and globin RNA along with ribosomal RNA was depleted from total blood RNA using Globin zero gold rRNA removal kit (Illumina cat.# GZG1224) to enrich poly-adenylated coding RNA as well as non-coding RNA. The rRNA and globin + rRNA depleted RNA from biopsy and blood total RNA, respectively was used for preparation of the sequencing library using RNA Tru Seq Kit supplied by Illumina (Cat # 1004814). The ribozero and globin zero RNA-Seq libraries were sequenced on the Illumina HiSeq 2500 platform using 100 bp paired end protocol following manufacturer’s procedure.

Genomic alignment to GRCh37 of single-end RNA-seq reads was performed using 2-pass STAR^2, 3^. Default parameters for STAR were used, as were those for the quantification of aligned reads to GRCh37.75 gene features via featureCounts^3^. Multimapping reads were flagged and discarded. Raw count data was pre-filtered to keep genes with CPM>0.5 for at least 3% of the samples. After filtering, count data was normalized via the weighted trimmed mean of M-values^4^.

Gene expression matrices were generated using the voom transformation and adjusted for technical variables (e.g. RIN, processing batch, rRNA rate and exonic rate) using the limma framework. Expression matrices were also adjusted for age, gender and genetic PCs for the GSVA analysis.  Statistical analysis was carried out using R language version 3.0.3^5^ and its available packages. Expression of ACE2 and TMPRSS2 were modeled using mixed-effect models with fixed factors depending on the comparison and a random intercept for each subject using the *nlme* package in R. Marginal means and hypothesis of interests were tested using the *emmeans* package capabilities. Effect of region and tissue (inflamed or non-inflamed) differences were estimated using a model that included disease (UC/CD/Non-IBD cohort), tissue and region and its interactions as well as age, gender and smoking status.

To investigate the medication effect on the MSCCR cross-sectional cohort, we first defined, for each medication, a propensity matched (PM) subcohort. The PM subcohort was defined such that patients not taking a given medication were selected to have the same distribution of clinical severity (SCCAI score for UC patients and HBI score for CD) and endoscopic disease severity (Mayo score for UC and SES-CD score for CD), number of surgeries, the availability of inflamed and non-inflamed biopsies at the time of endoscopy than those taking the medication. For each medication, data from the PM subcohort was modeled using a linear mixed-effect model with fixed factors subtype (CD, UC), medication, region, tissue and its interactions.

**Curation of RNA-seq based molecular signatures related to IBD and COVID-19 response**

*We curated RNA-seq based molecular signatures related to IBD and COVID-19 response by identifying differentially expressed genes (DEGs) as*

a) IBD: DEGs between IBD inflamed and non-inflamed biopsies (FDR<0.05 and fold change>2) compared using a mixed-effect model with a random intercept for each MSCCR patient and fixed factors for tissue, region and disease.
b) SARS-CoV-2-*infection of epithelia:* Genes associated with COVID-19 response in two epithelial model systems recently published^6, 7^, comparing SARS-CoV-2 infection of:
   i. i) A549 or **NHBE COVID-19:** SARS-CoV-2 infection of: 1. A549 lung alveolar carcinoma cells or 2. normal human bronchial epithelium (NHBE) for 24 hours^7^ (FDR <0.05)
   ii. ii) DIF or EXP **hSIOs COVID-19:** human small intestinal organoids (hSIOs) grown in either1.Wnt high expansion (EXP) medium or 2. differentiation (DIF) medium^6^ (FDR <0.1 or <0.05).
c) *Blood in* SARS-CoV-2-*infection:* DEGs in whole blood RNAseq profiles of 76 adult patients with SARS-CoV-2 pneumonia versus 24 healthy controls (FDR <0.05 and |FC|>1.5) as part of the Hellenic Sepsis Study group^8^.
d) *IBD drug response signatures*: We analyzed publicly available gene expression profiles from UC and CD patients treated with infliximab (GSE16879)^9^ and CD patients treated with ustekinumab (GSE112366)^10^. We defined treatment and tissue specific DEGs by comparing gene expression after treatment course to baseline expression in patients who responded to treatment.
   i. i) Ileum CD infliximab or ustekinumab response: Gene signatures representing the response to infliximab (GSE16879)^9^ and ustekinumab (GSE112366)^10^ in ileal biopsies of CD patients were generated (at unadj P<0.01).
   ii. ii) Colon UC infliximab response: Infliximab response-associated gene signatures in colonic biopsies from a cohort with UC (FDR<0.01) that were responding at week 4-6 GSE73661^11^).
   iii. iii) Colon CD infliximab or ustekinumab response: Infliximab response-associated gene signatures in colonic biopsies from a cohort with CD that were responding at week 4-6 weeks (GSE16879)^9^FC>|1.5|, FDR<0.05) were generated. Ustekinumab response signature in the colon of patients with CD was generated in colonic biopsies of week 6 responders vs baseline (FC>|2| and FDR <0.05)^12^.

**Analyses of the CERTIFI Cohort**

Eighty anti-TNFα refractory CD patients enrolled in a phase 2b crossover trial (CERTIFI trial)^13^ were randomized to ustekinumab or placebo at baseline and received the assigned treatment until week 8 (induction period)^12^. Clinical response at week 22 was defined as a decrease of 100 or more in Crohn’s Disease Activity Index (CDAI) score from baseline. Microarray (HT_HG-U133_Plus_PM) gene expression data (available at GSE100833) from 810 biopsy samples taken at baseline, weeks 6 and 22 were obtained for further analysis. The effect of treatment during the induction period (574 biopsies from 80 patients) was modeled using a linear mixed-effect model with visit, tissue, region, treatment group and its interactions. Changes over time for each treatment/region/tissue were tested using the *emmeans* package. Differences between week 22 responders and non-responders were evaluated only in endoscopically defined inflamed samples (n=108) from patients (n=28) who were always on ustekinumab using a mixed-effect model with visit, region and response as fixed effect and its interactions. Gene expression changes over time (screening, week6, week22) were estimated for responders and non-responders.  For 227 patients, blood transcriptome data was available and a similar analysis strategy was used.

**Analyses of GEMINI I and GEMINI LTS**

The GSE73661 series included expression profiles from moderate-to-severe UC patients enrolled in GEMINI-I and GEMINI LTS trials evaluated the efficacy of vedolizumab^11^. Colonic biopsies were obtained from 44 (41 receiving vedolizumab and 3 receiving placebo) patients at baseline and week 6. Response was defined as endoscopic mucosal healing (Mayo endoscopic subscore 0 or 1) at week 6. This series also included colonic biopsies from 23 UC patients treated with infliximab at baseline and after 4-6 weeks of treatment; the same definition of endoscopic mucosal healing was used. The treatment effect after the induction period (4-6 weeks) was estimated using a mixed-effect model with time and treatment group and its interactions as fixed effects. Changes between endoscopic responders and non-responders were evaluated on a similar model including the interaction of time and response.

**System Biology approach integrating IBD Bayesian Networks and COVID signatures:**

**MSCCR genotype generation:**

DNA was isolated from whole blood using QIAamp®DNA BloodMini Kit (QiagenCatalogNo.51104). Genotype data generated using the high-density Illumina Multi-Ethnic Global Array (MEGA^EX^) and Infinium ImmunoArray-24 v2 BeadChip arrays. We further imputed genotypes using the Michigan Imputation Server^9^ with the 1000 Genomes reference.  QC filtering was performed within each batch and filtering criteria were: missing rate per sample and per probe ≤ 10%; HWE p-value per probe > 1E-6; identity between self-reported and inferred sex; samples pairwise Identity By Descent PI_HAT < 0.8; visual identification of samples outliers within the first 2 PCs. In addition, consistency of technical replicates was checked.

**MSCCR Bayesian gene regulatory network (BGRN) generation:**

Bayesian gene regulatory networks (BGRNs) can capture fundamental properties of complex systems in states that give rise to complex (diseased) phenotypes^12^. We and others have successfully identified and validated a large number of novel targets using these derived network models. Such approaches have greatly expanded our understanding of complex diseases such as diabetes, Alzheimer’s and IBD^12, 14, 15^. Bayesian networks were generated from RNA sequence data generated on intestinal biopsies from the MSCCR cohort using their intestinal expression QTL information (eQTLs) as priors.  The Bayesian networks were region- (ileum or colon/rectum) and disease- (CD, UC, and control) specific and included both inflamed and uninflamed biopsies.  MSCCR Bayesian networks were reconstructed using RIMBAnet software^16-18^ as previously described and visualized using Cytoscape 3.7^19^. RIMBAnet software is available with step by step instructions. Final networks were decided with Monte Carlo Markov Chain (MCMC) simulation^20^ which creates thousands of possible different networks, which were then combined to form a consensus network.  We also used two publicly available Bayesian networks. One was built from ileum biopsy data collected from treatment naïve Crohn Disease pediatric patients (RISK cohort) as previously described^12, 21^.  The second was generated using data from the CERTIFI cohort which included anti-TNFα refractory CD patients from whom biopsies taken across multiple intestinal regions (ileum, ascending colon, descending colon, sigmoid colon, and rectum inflamed and non-inflamed tissue)^12^.

**Bayesian subnetwork generation:**

*ACE2 and TMPRSS2 subnetwork*: Gene-centric subnetworks were generated by selecting either ACE2 or TMPRSS2 on various Bayesian networks (Ileum CD, Pancolonic CD, and Pancolonic UC) and expanding out three to five layers (undirected) to obtain the nearest ACE2 or TMPRSS2 neighbors.  The connected subnetworks obtained were generally between 200-500 genes in total.

*Differential gene expression signatures related to*:

a) *IBD inflammation*: MSCCR IBD inflammation genes were defined as differentially expressed genes (FDR<0.05 and fold change>2) between IBD inflamed and non-inflamed samples contrasted in a mixed-effect model with a random intercept for each patient and fixed factors for tissue, region and disease as well as core technical and demographic covariates.

b)	SARS-CoV-2-*infection of epithelia:* We curated molecular signatures associated with COVID-19 response in various epithelial model systems recently published^6, 7^. SARS-CoV-2 infection of: i) A549 lung alveolar carcinoma cells or normal human bronchial epithelium (NHBE) for 24 hours^7^ (FDR <0.05) or ii) human small intestinal organoids (hSIOs) grown in either Wnt high expansion (EXP) medium (at FDR<0.1 or <0.01) or differentiation (DIF)^6^ medium (at FDR <0.1 or <0.05). To have comparable signature set sizes differential expression significance, depending on the analysis, for DIF-hSIO was defined as either FDR <0.1 or 0.05 and either FDR <0.05 or 0.01 for the EXP-hSIO model. A signature representing commonly up-regulated lung and gut model COVID-19 responsive genes was also derived by first performing a union of up-regulated genes within either the lung (at FDR<0.05 =443 genes) or gut (at FDR<0.1 = 283) models and then intersecting the two-tissue model genesets (49 up-regulated genes in common).

b) *Blood in* SARS-CoV-2-*infection:* We curated a whole blood RNAseq signature which was generated on 76 adult patients with SARS-CoV-2 pneumonia and 24 healthy controls as part of the Hellenic Sepsis Study group^8^ at FDR <0.05 and threshold of > |1.5|.
c) *IBD drug response-associated gene expression signatures*:

Ileum: Gene signatures representing the response to infliximab (GSE16879)^9^ and ustekinumab (GSE112366)^10^ in ileal biopsies of CD patients were generated (at unadj P<0.01). The GSE16879^9^ series was used to generate a differential expression signature associated with infliximab response in CD patients. The patients were classified for response to infliximab based on endoscopic and histologic findings at 4-6 weeks after first infliximab treatment. Ileal biopsies in responders were compared to baseline samples using the *limma* framework. For the ustekinumab signature, the series GSE112366^10^ including microarray expression profiles from biopsies of moderate-to-severe CD patients enrolled in two phase 3 studies (UNITI-2 and IM-UNITI) were used. These patients failed conventional therapies previously and were largely naive to anti-TNFα therapy. Ileal biopsies from responders (based on mucosal healing) at 8 weeks were compared to baseline using the *limma* framework to identify ustekinumab response genes.

Colon: Infliximab response-associated gene signatures were generated from colonic biopsies sampled from a cohort with UC (FDR<0.01 at week 4-6 responders compared to week 0) (GSE73661)^11^ and a cohort with CD (FC>|1.5|, FDR<0.05) (GSE16879)^9^. An ustekinumab response signature in the colon of patients with CD^12^ was generated comparing biopsies from week 6 responders to baseline (FC>|2| and FDR <0.05).

IBD Inflammation, COVID-19 and IBD Drug Response- Subnetwork generation:

Genes found altered in NHBE/A549 or organoids following SARS-CoV-2 infection; IBD inflammation; or response to medications were separately projected onto various Bayesian networks (Ileum CD, Pancolonic CD, Pancolonic UC) allowing 1 or 2 nearest neighbor to be included. The most connected subnetworks were then extracted to generate model-specific SARS-CoV-2 infection-; IBD inflammation-; or drug-response associated subnetworks (see Supplementary Methods Table 1 below).  The genes common between these networks were determined and tested for enrichment to various genesets using the Fisher’s exact test and p-values were adjusted using Benjamini-Hochberg procedure.
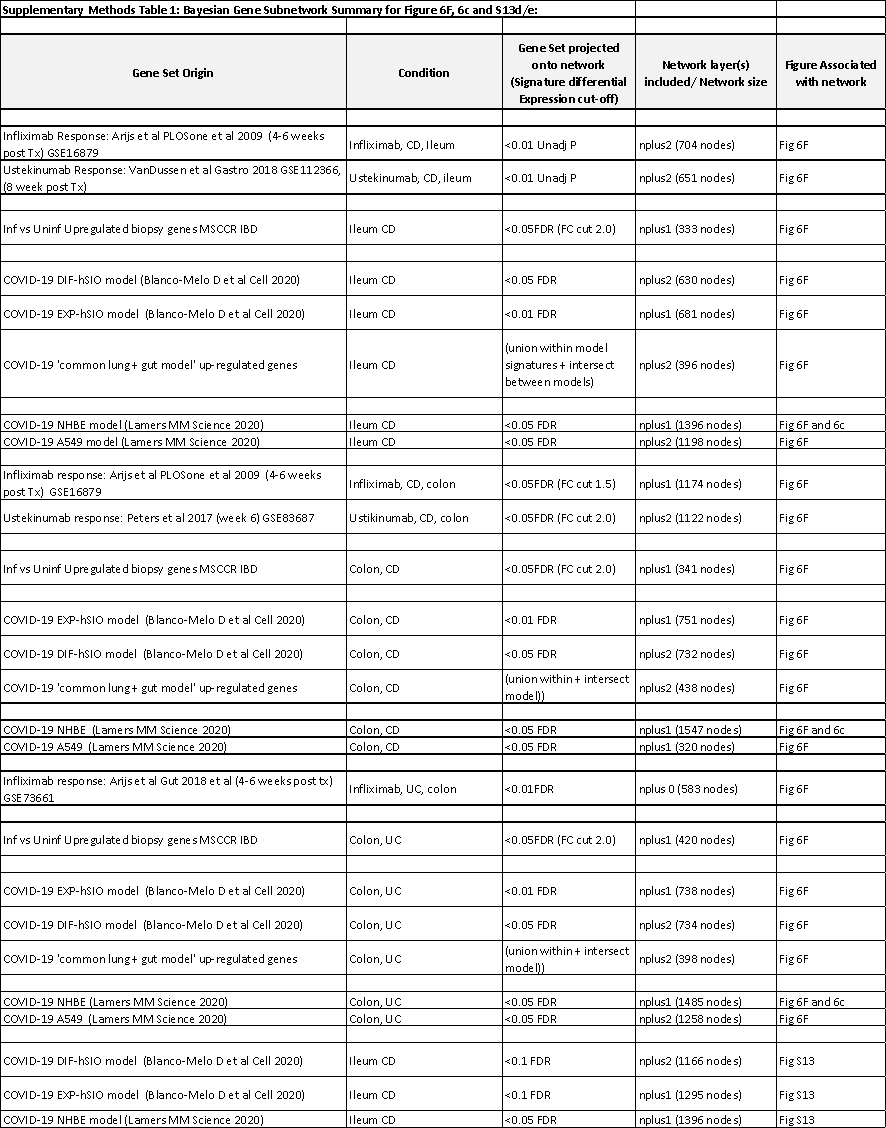

**Pathway and geneset enrichment analysis of subnetworks:**

Gene subnetworks were tested for functional enrichment using the Reactome pathway database.  Reactome pathway gene sets sourced from Enrichr^22^ and tested for enrichment using a Fisher’s exact test with Benjamini- Hochberg multiple test correction. ACE2 and TMPRSS2 associated subnetworks were also tested for enrichment in gut cell types using genesets from single RNA cell data from Smillie et al^23^ and Huang et al^24^. Enrichment in genesets associated with various macrophage perturbations (e.g. cytokines)^25^, and ACE2 co-expressed genes^26^ as well as reported IBD GWAS genes^27-29^ were also tested.

**Key driver gene analysis:**

Key driver analysis (KDA) identifies key or “master” driver genes for a given gene set in a given gene regulatory network.  We used a previously described KDA algorithm by Zhang et al^30^. KDA requires two input files, a set of genes (G) and a (un)directed gene network (N). A subnetwork N_g_ is first defined as the set of nodes in N that are no more than h-layers away from the nodes in G. For this analysis we used the whole network as Ng. We then used a dynamic neighborhood search mode (DNS) feature which searches the h-layer neighborhood (HLN, h=3) for each gene in Ng (HLNg,h) for the optimal h* giving the maximum computed enrichment statistic for HLN(g,h). A node becomes a key driver, if its HLN is significantly enriched for the nodes in G (at adj P <0.05). The set of genes for KDA included either the NHBE COVID infection geneset or the IBD inflammation geneset. Key driver genes were further summarized by frequency in which they appeared across all networks.

**Geneset variation analysis of SARS-CoV-2 infection gene expression signatures:**

We employed the blood and epithelial model COVID-19 response gene signatures in a gene set variation analysis (GSVA) using MSCCR biopsy or blood RNAseq data and CERTIFI blood and biopsy microarray data. For each gene set (up or down-regulated), GSVA, a non-parametric and non-supervised method, estimated the overall variation of the gene set on the expression profiles of the MSCCR biopsy and blood expression matrix after adjusting for gender, age, genetic PCs and technical covariates. As a result, a z-score like sample-wise enrichment score was calculated for each gene set. Such enrichment scores were then used for hypothesis testing with respect to phenotype information.

**T cell transfer model:**

We compiled temporal gene expression profiles that were recently published throughout the development of CD4CD45Rbhi T cell transfer colitis. Fang et al^31^ performed the CD4CD45Rbhi T cell transfer colitis model and colonic tissue was used for genome expression profiling analysis at 0, 2, 4, or 6 weeks after adoptive T cell transfer. 1775 genes were identified as differential expressed during the progression of T cell mediated colitis, and they classified these genes according to 8 temporal phenotypes. We utilized two temporal phenotype patterns: ‘W0’ which were genes progressively downregulated over the 6 weeks (aka W0>W2>W4>W6) and ‘W6’ which were genes found progressively upregulated over the 6 weeks (aka W0<W2<W4<W6). Genes were converted to human symbols for analysis.

**DSS mouse model:**

The dextran sodium sulfate (DSS) mouse model commonly employs a 5 to 7 day DSS exposure after which a colitis-like macroscopic phenotype is observed. A recent paper compared molecular responses of a DSS model during the colonic inflammation phase (during DSS treatment) followed by tissue regeneration phase (every 2 days post DSS treatment up to day 14) to a differential expression signature comparing UC vs healthy patient biopsies. The authors found ~650 genes in common^32^. We therefore utilized this signature which supports molecular parallels between the DSS mouse model of IBD and adult IBD in order to compare to the COVID-19 response signatures.

**TNBS mouse model:**

We curated gene expression signatures from a TNBS associated experiment (FCH>0.5 and adj.P-value <0.05) which involved evaluating the recurrent molecular responses in the colons by giving three weekly intra-rectal instillations of TNBS allowing for both acute (active inflammation) and chronic processes of IBD to be assessed as sampling was done before and 2 or 7 days after each TNBS instillation^31^. On days 7, 14 and 21 mice were administered intra-rectally TNBS, at selected time-points 2 and 7 days after each TNBS administration (ie Day 9, 14, 16, 21, 23 and 28), mice were sacrificed and molecularly profiled. The 6 resulting genesets included: 1. Day 9 = (2 days Post TNBS administration); 2. Day14 = (Before second TNBS administration); 3. Day16 = (2 days post second TNBS administration)’ 4. Day21 = (Before third TNBS administration); 5. Day23 = (2 days post third TNBS administration) and 6. Day28 = (7 days post third TNBS administration)

**URLS:**

RIMBANET: https://labs.icahn.mssm.edu/zhulab/?s=rimbanet&submit=Search
