## Supplementary figures and images for "Intestinal inflammation modulates the expression of ACE2 and TMPRSS2 and potentially overlaps with the pathogenesis of SARS-CoV-2 related disease"

### Supplementary Figure 2

Supplemental Figure 2

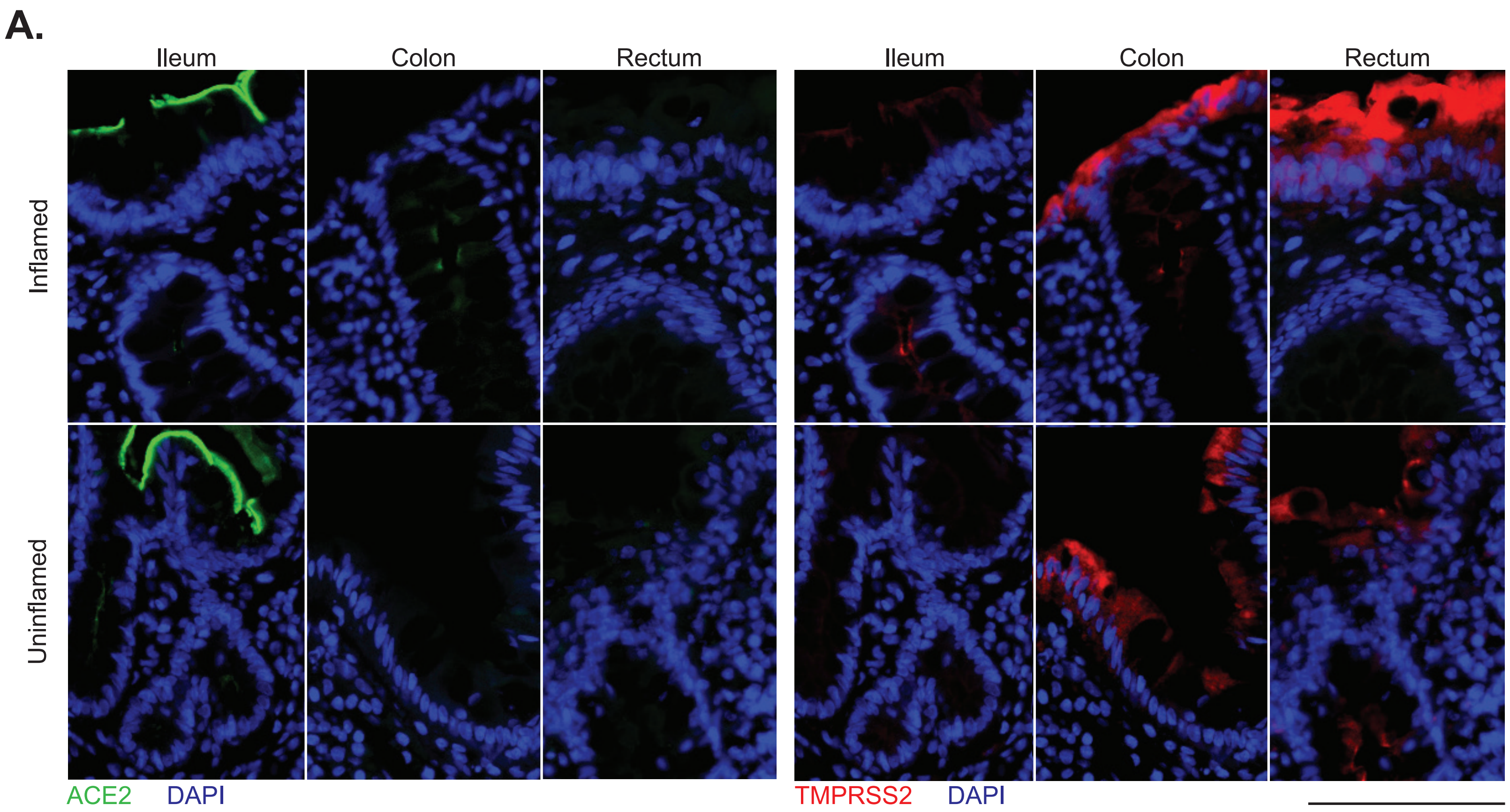

### Supplementary Figure 3

Supplementary Figure 3

a. ACE2

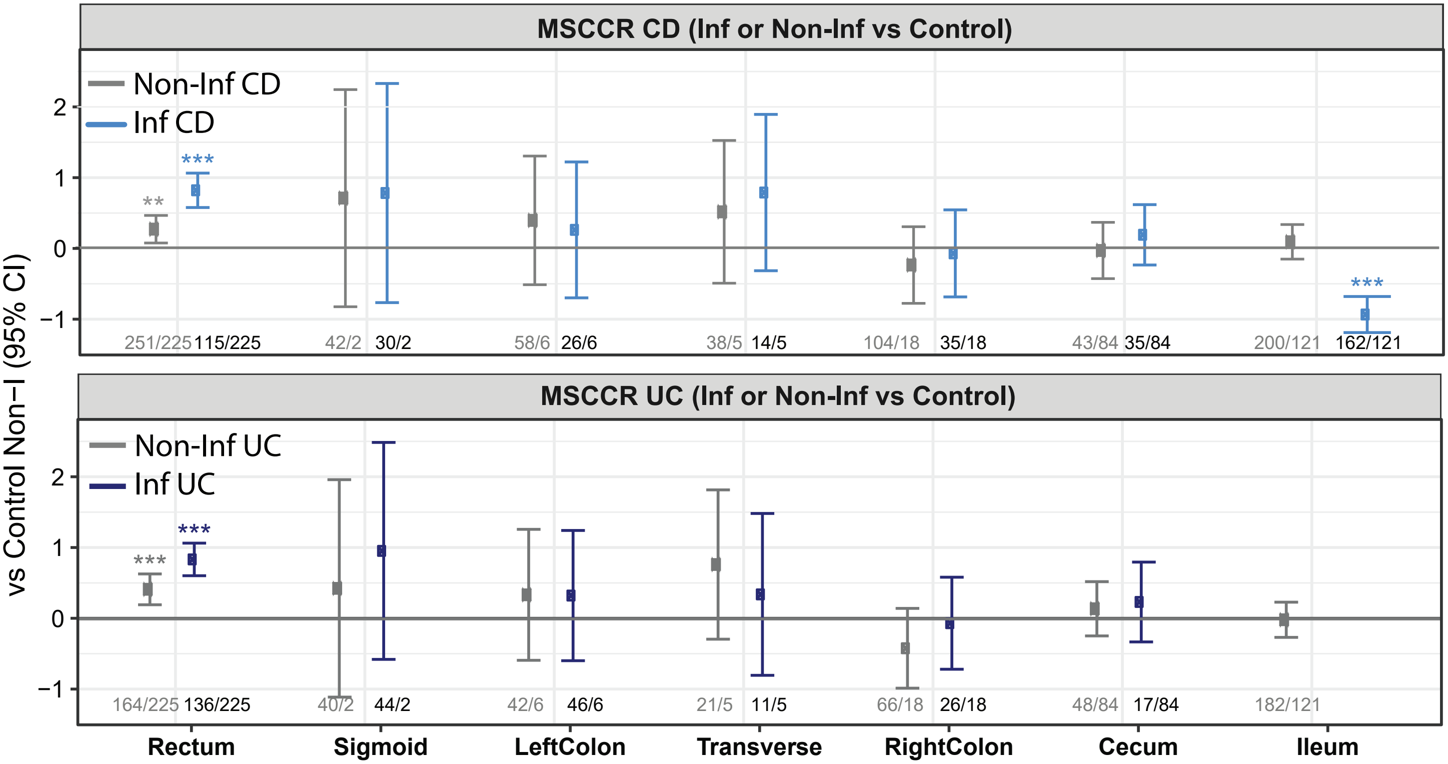


b. TMPRSS2

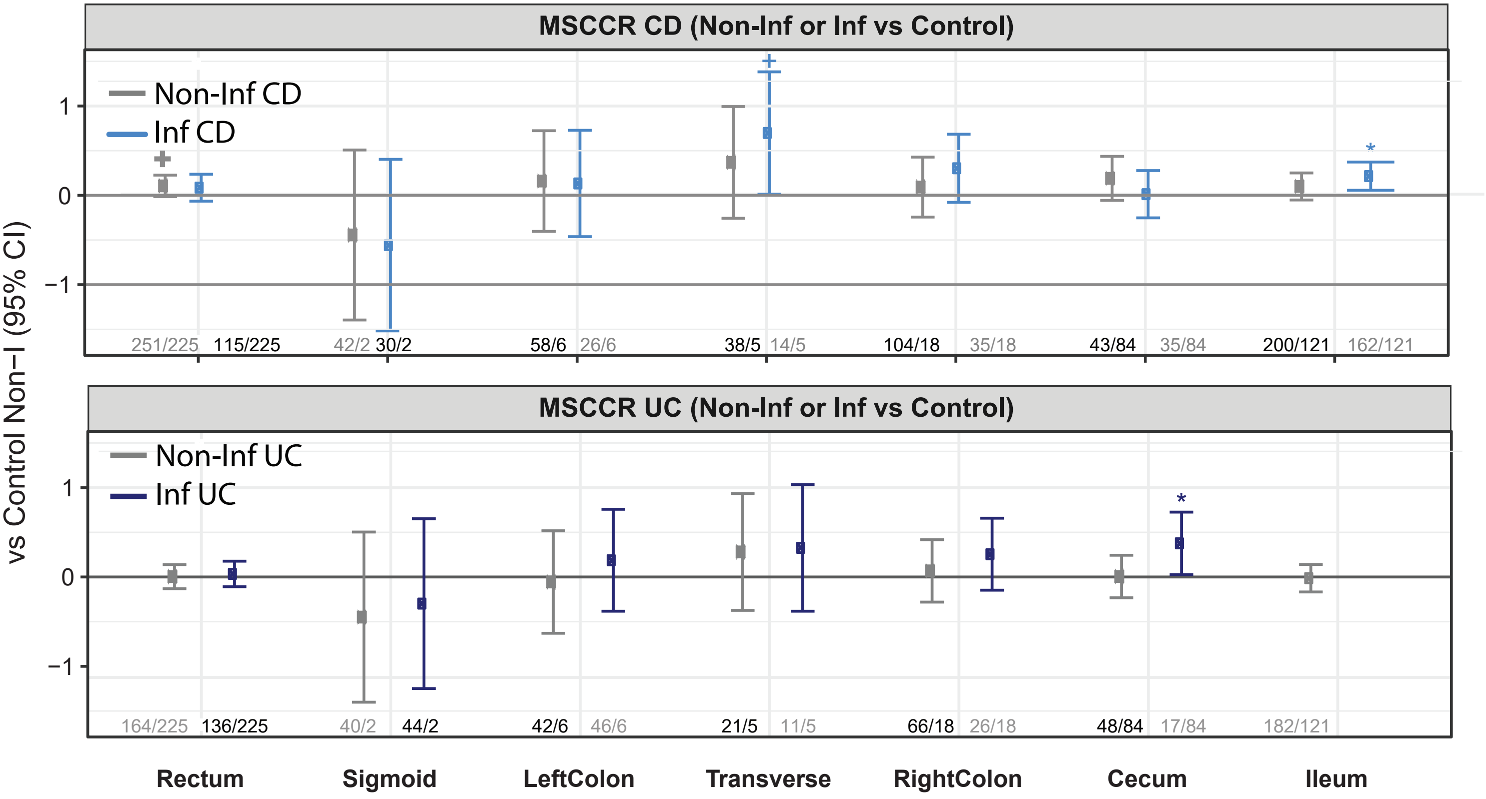

### Supplementary Figure 4

Supplementary Figure 4

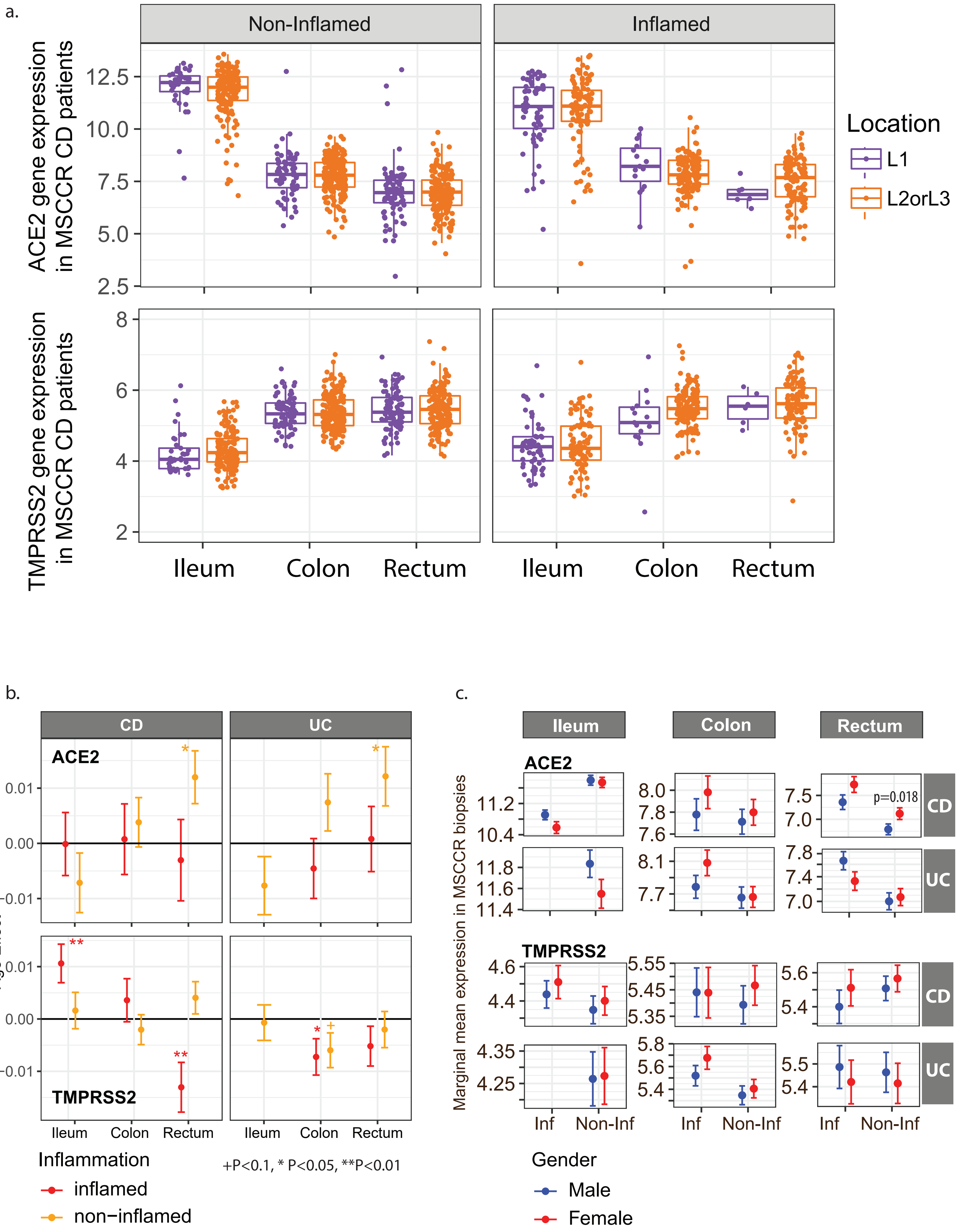

### Supplementary Figure 5

Supplementary Figure 5

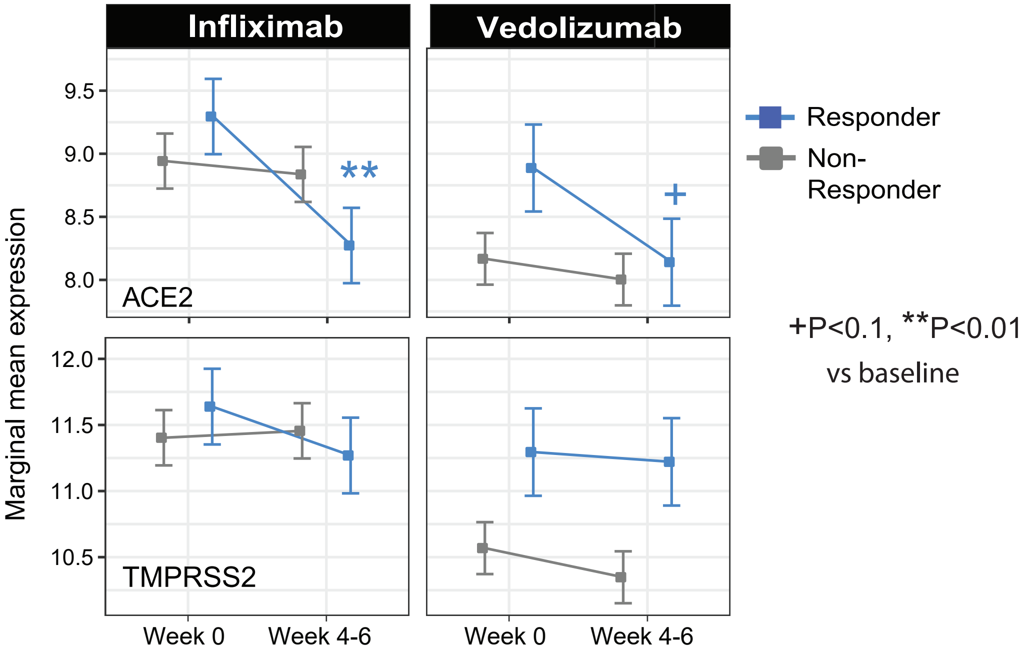

### Supplementary Figure 6

Supplementary Figure 6

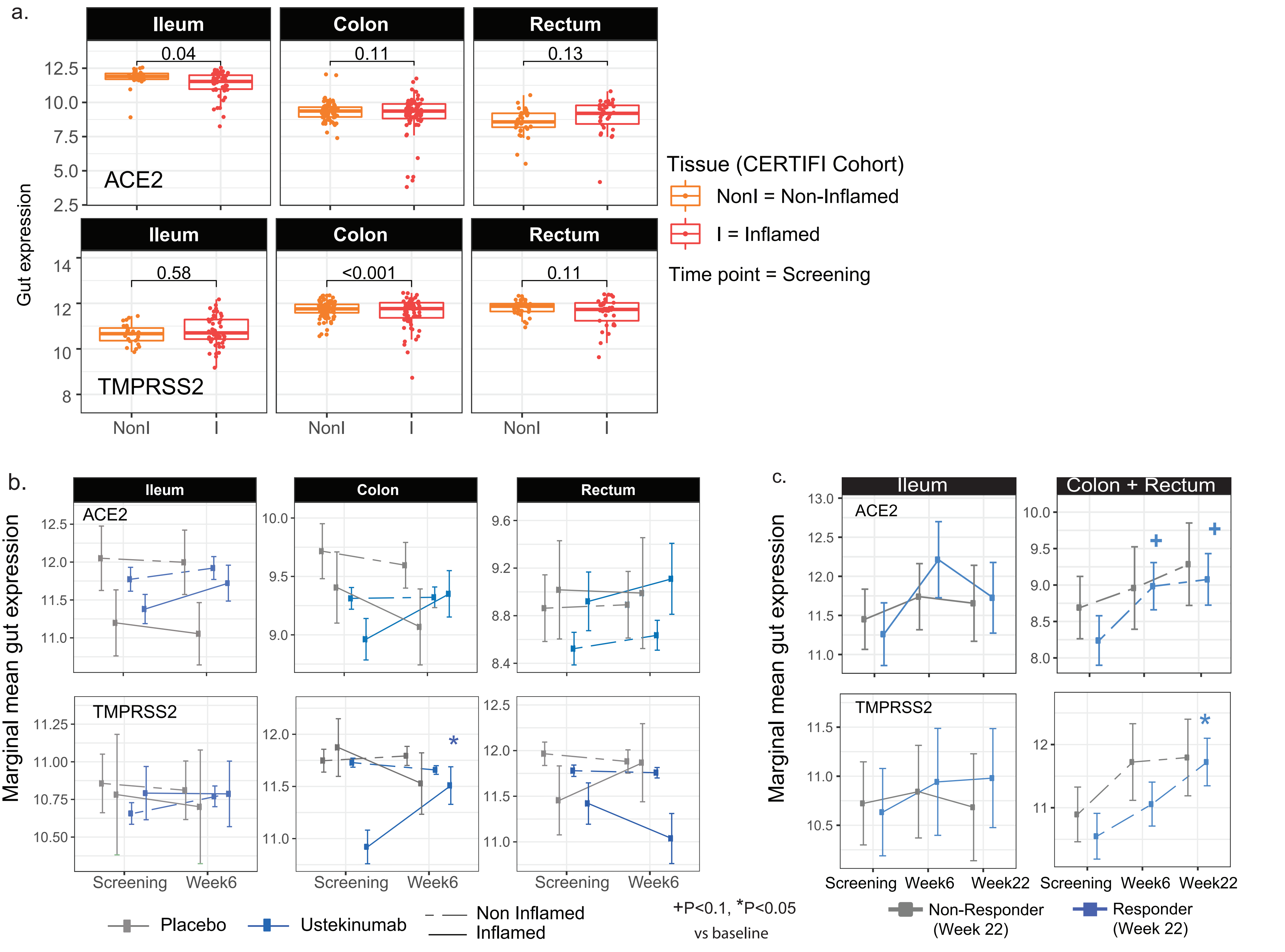

### Supplementary Figure 7

Supplementary Figure 7

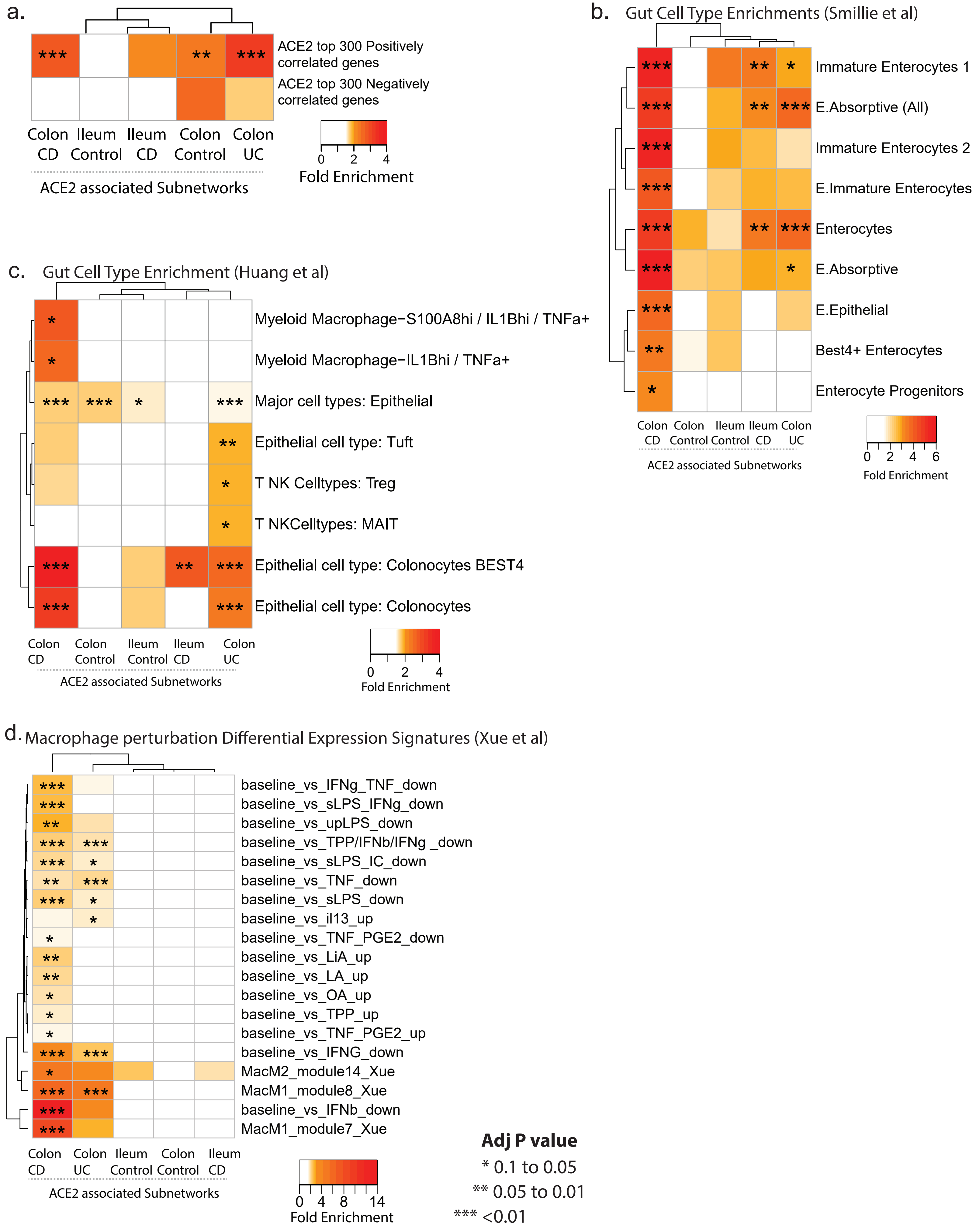

### Supplementary Figure 9

#### Supplementary Figure 9

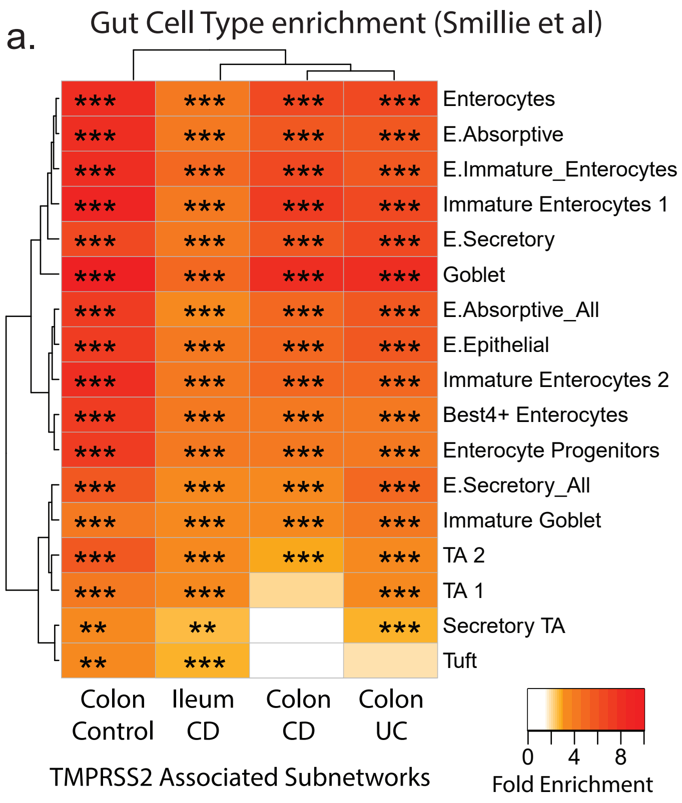


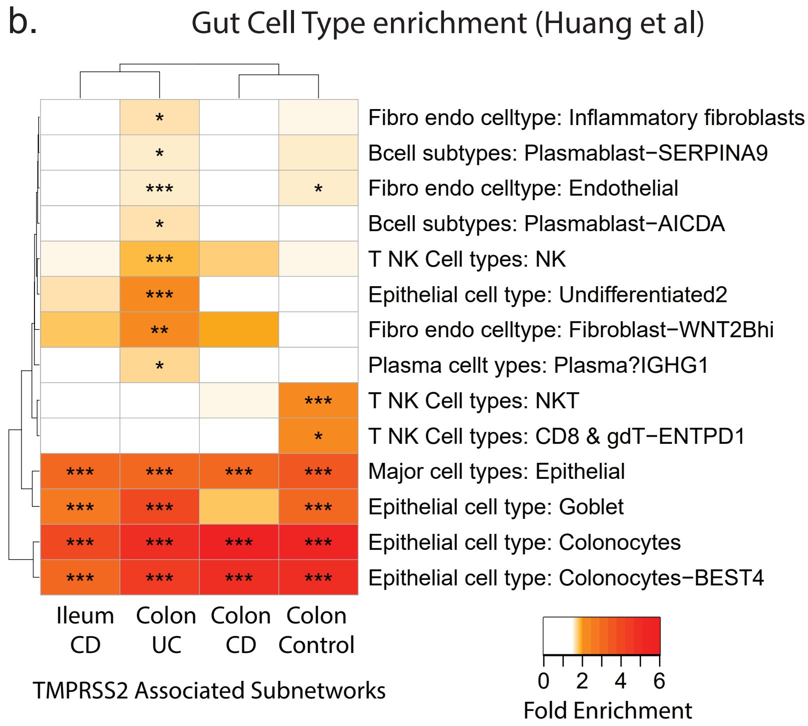


Adj P value: * = 0.1 -0.05; ** = 0.05-0.01; *** <0.01

### Supplementary Figure 10

Supplementary Figure 10

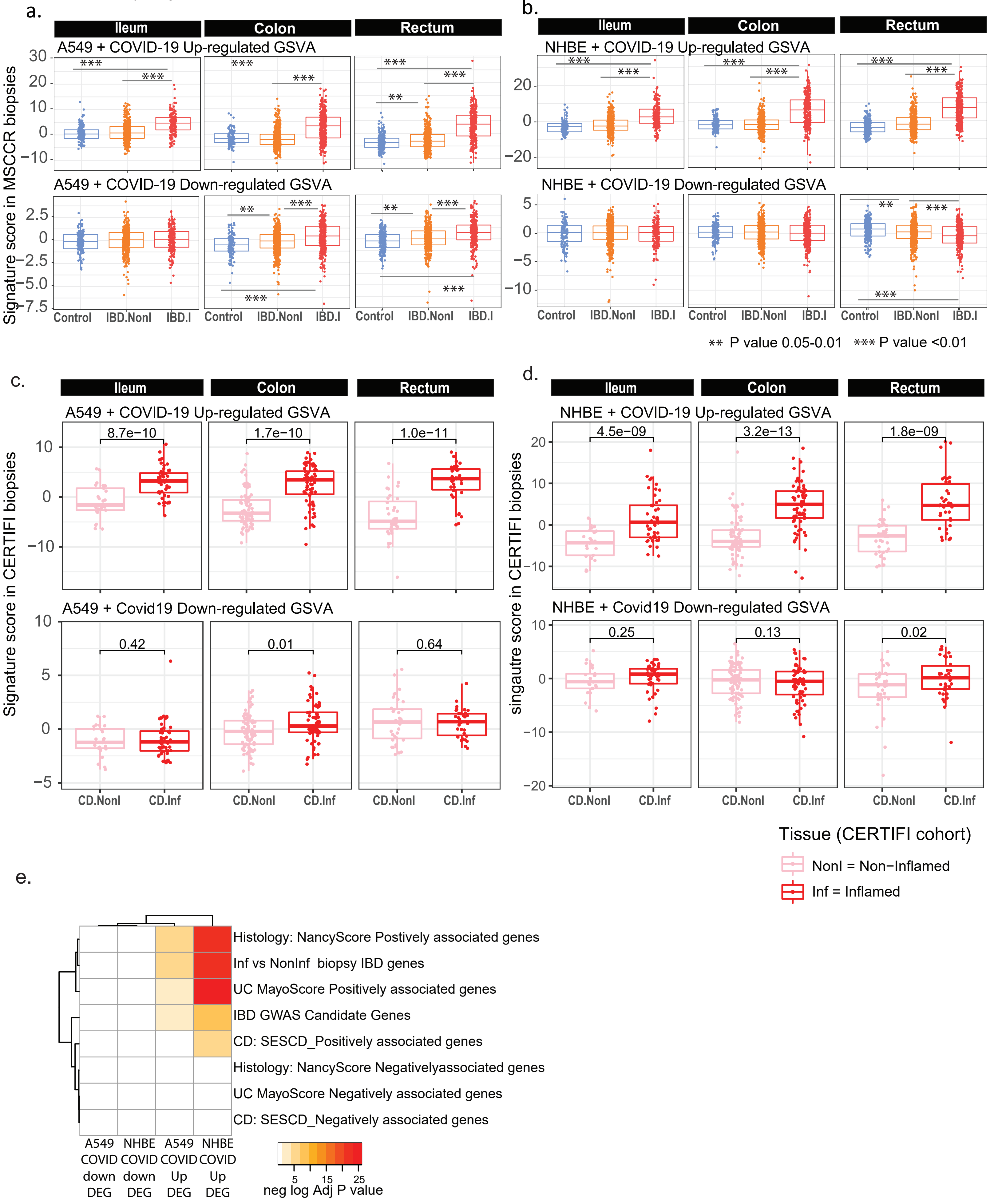

### Supplementary Figure 11

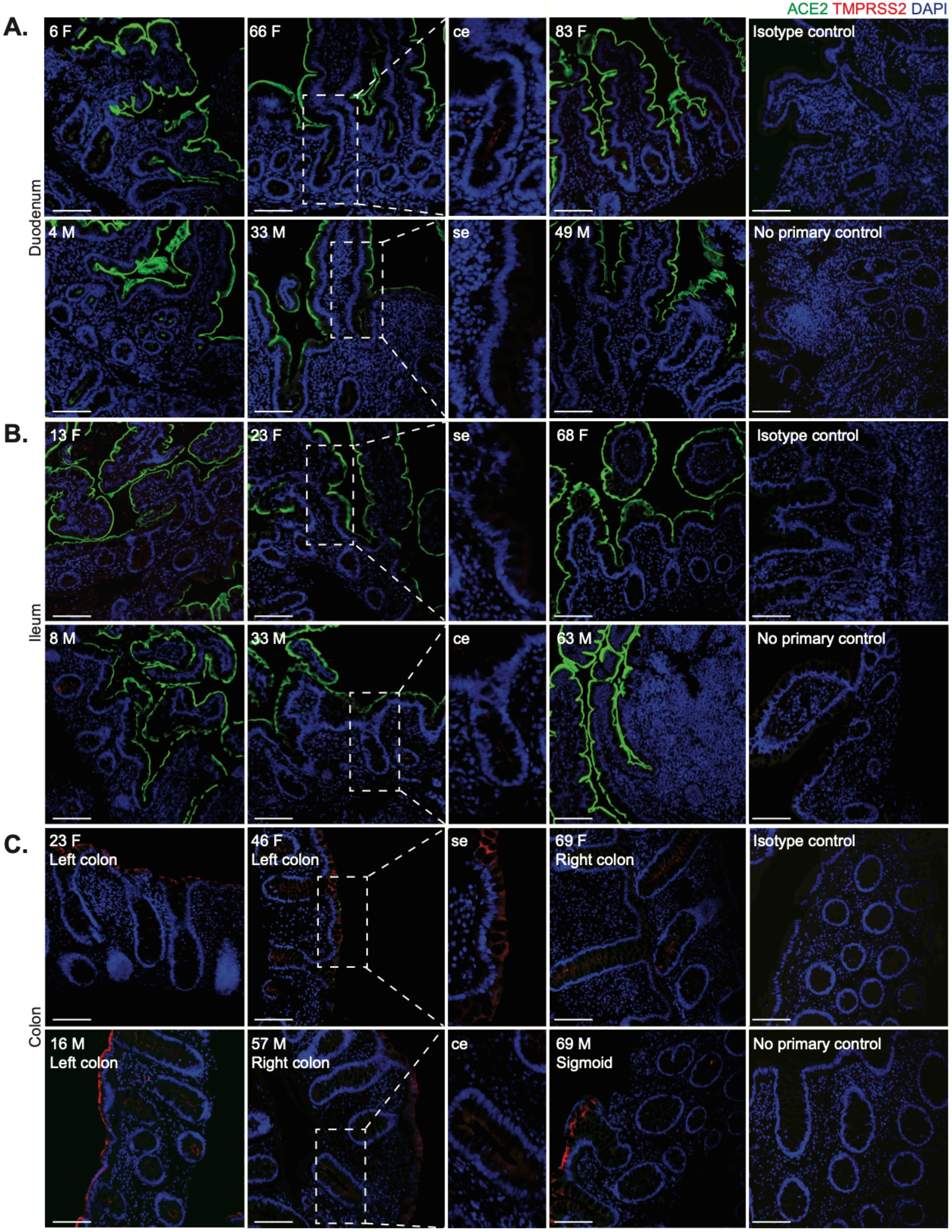

### Supplementary Figure 11

Supplementary Figure 11

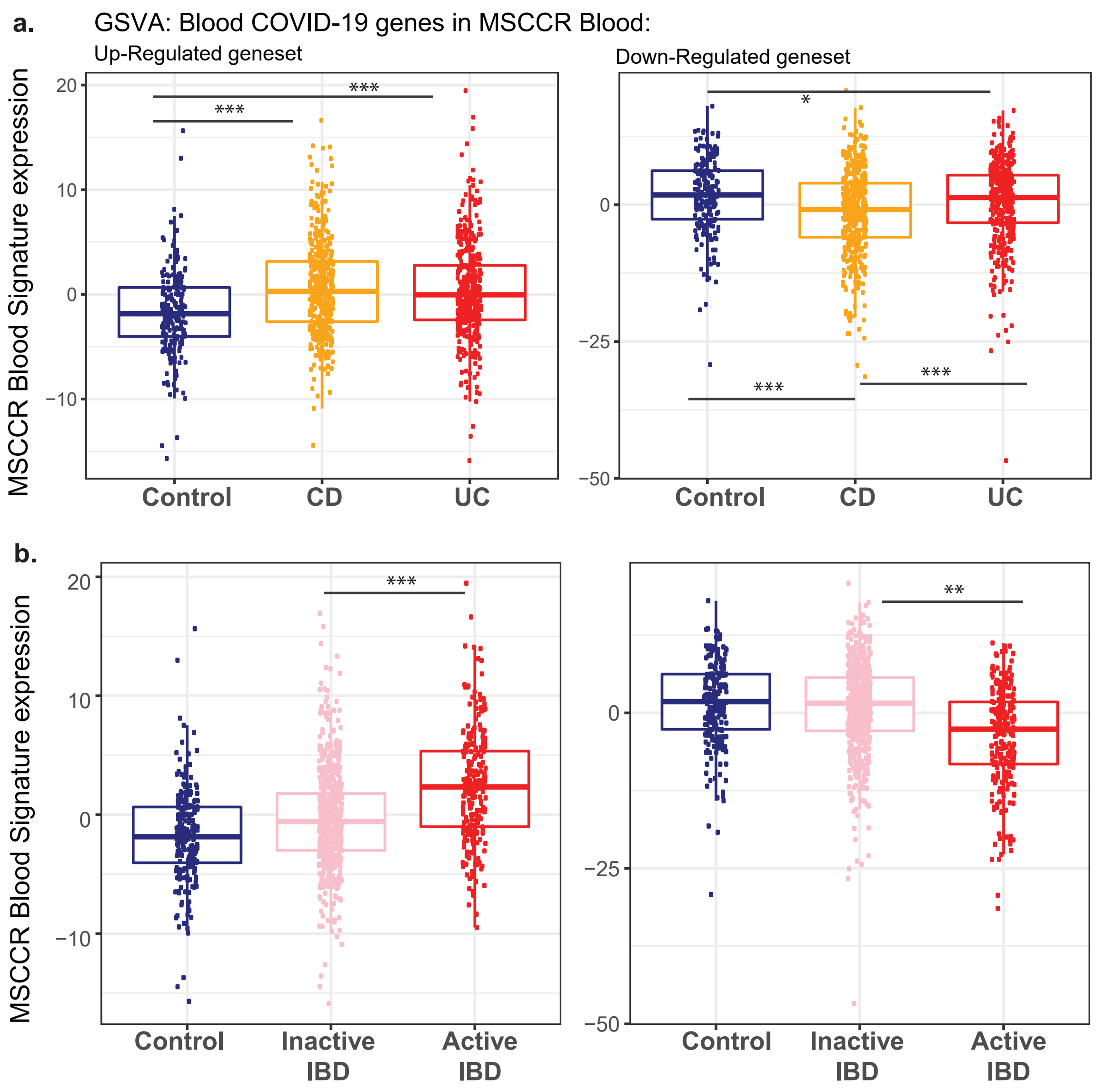

### Supplementary Figure 12

Supplementary Figure S12

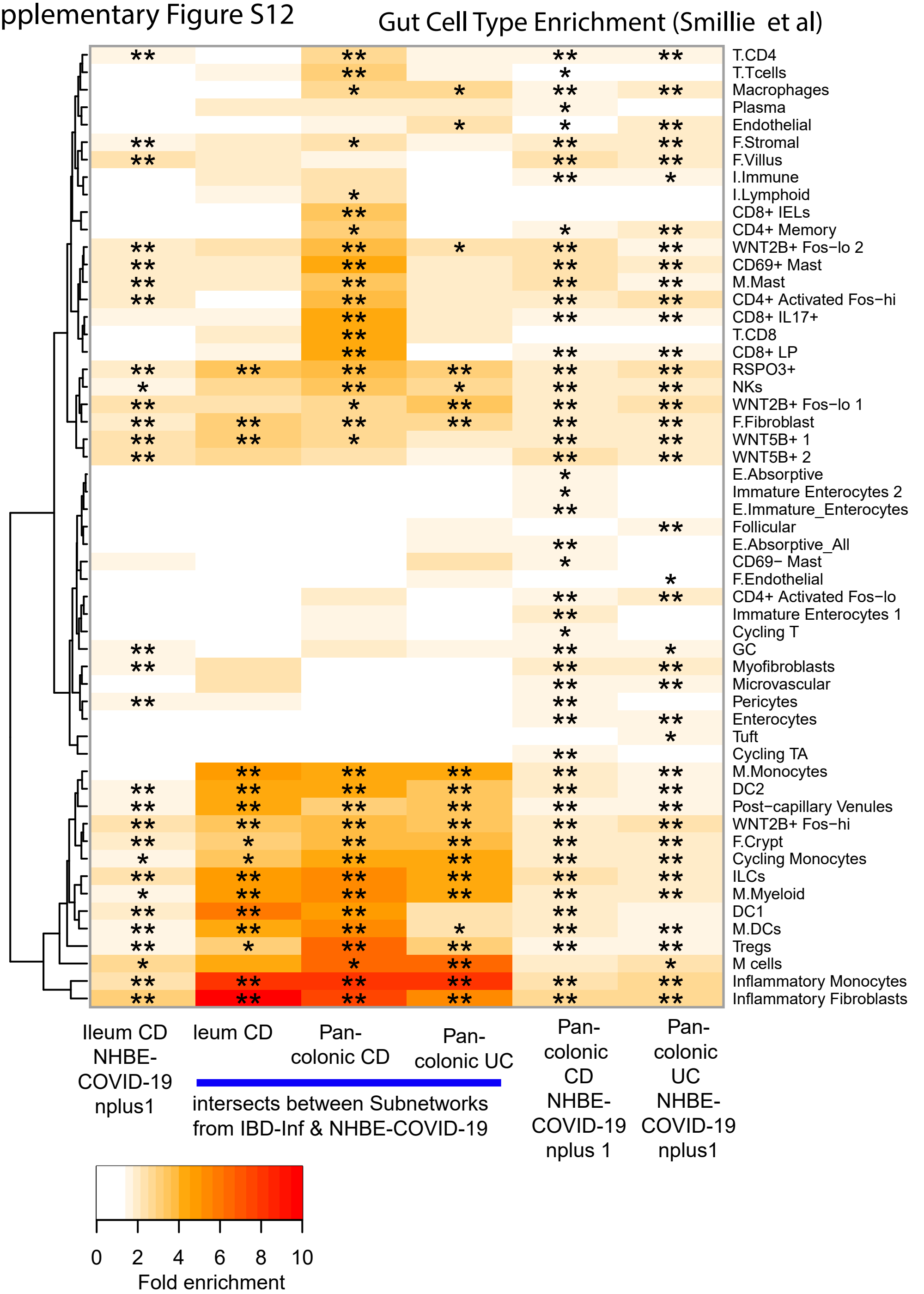


Adj P value * <0.05 ; *<0.01; ***<0.01

### Supplementary Figure 13

Supplementary Figure 13

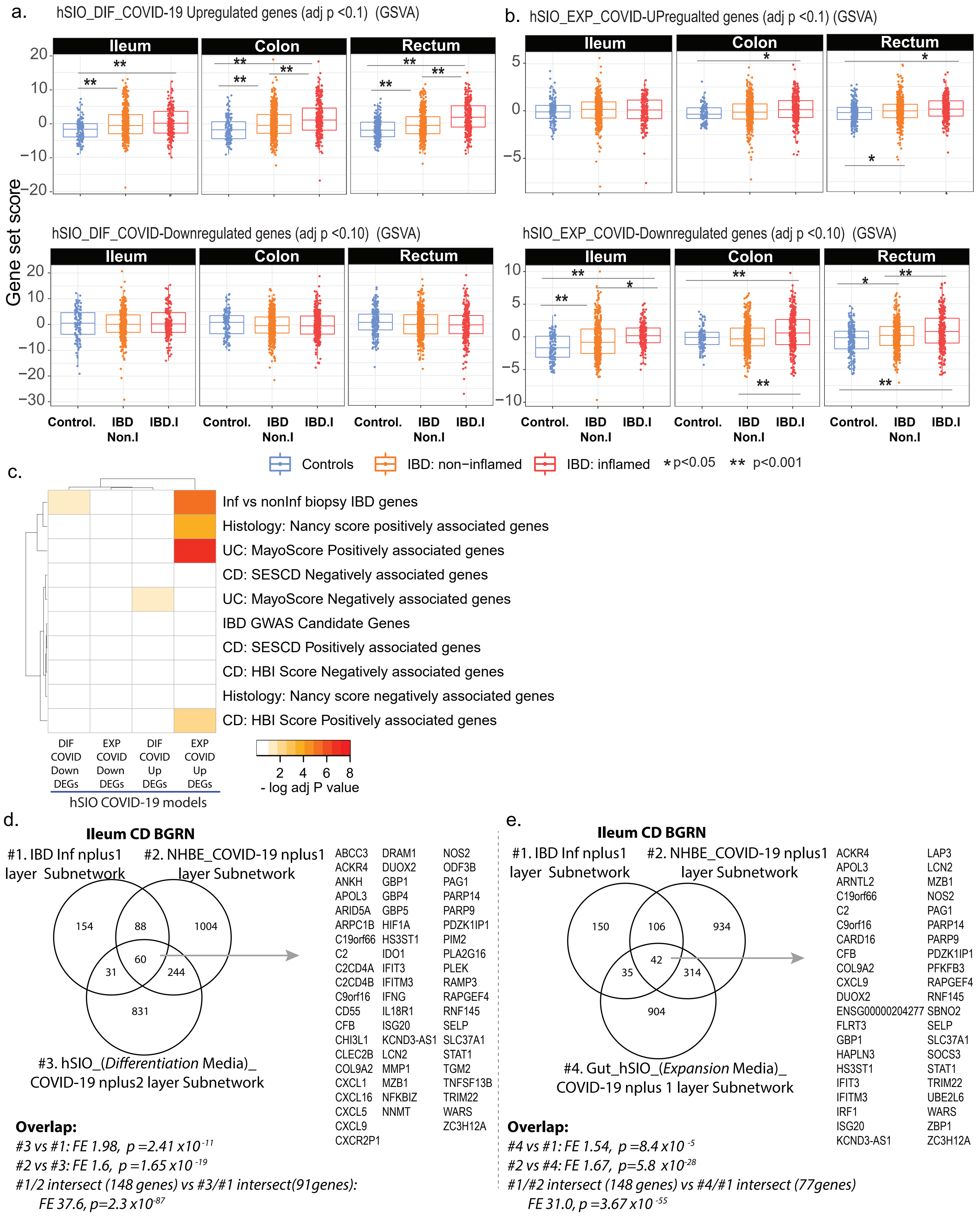

### Supplementary Figure 14

Supplementary Figure S14

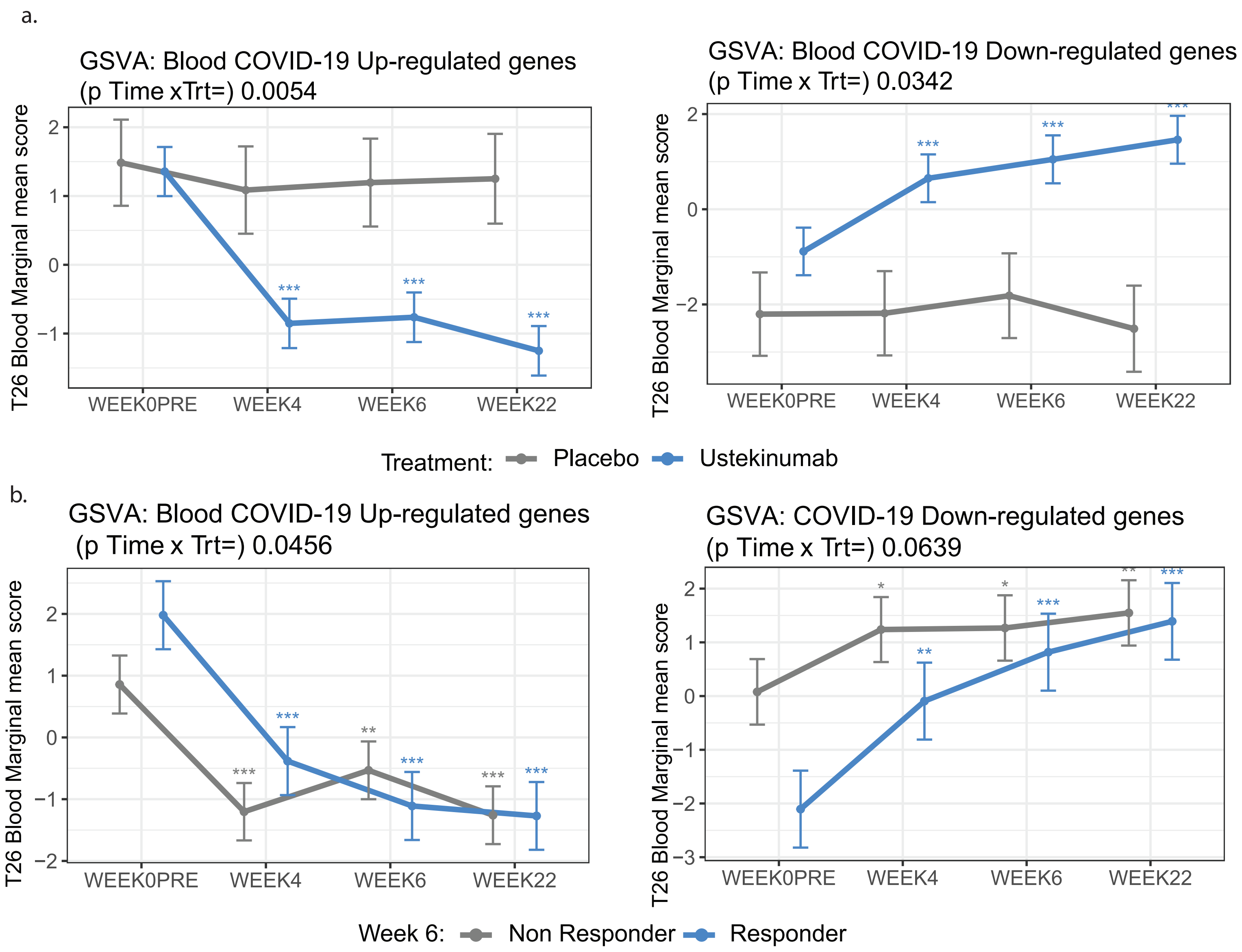
