## Supplementary Figure 8 for "Intestinal inflammation modulates the expression of ACE2 and TMPRSS2 and potentially overlaps with the pathogenesis of SARS-CoV-2 related disease"

a. Colon UC BGRN

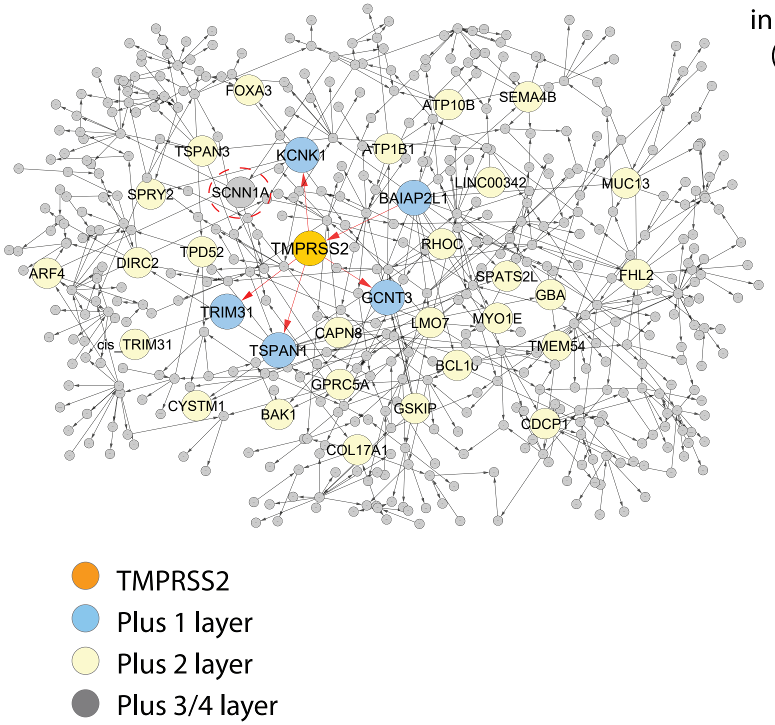

b. TMPRSS2 associated genes in 4 of 4 subnetworks (1 ileum + 3 colon)

| |
|---|
| BAIAP2L1 |
| BCL10 |
| CAPN5 |
| DSC2 |
| ELF3 |
| EPHA2 |
| F11R |
| GCNT3 |
| GPD2 |
| GSKIP |
| HN1 |
| MYO1E |
| OAS1 |
| RIOK3 |
| SLC44A4 |
| SSFA2 |
| TMPRSS2 |
| TRIM31 |

c.

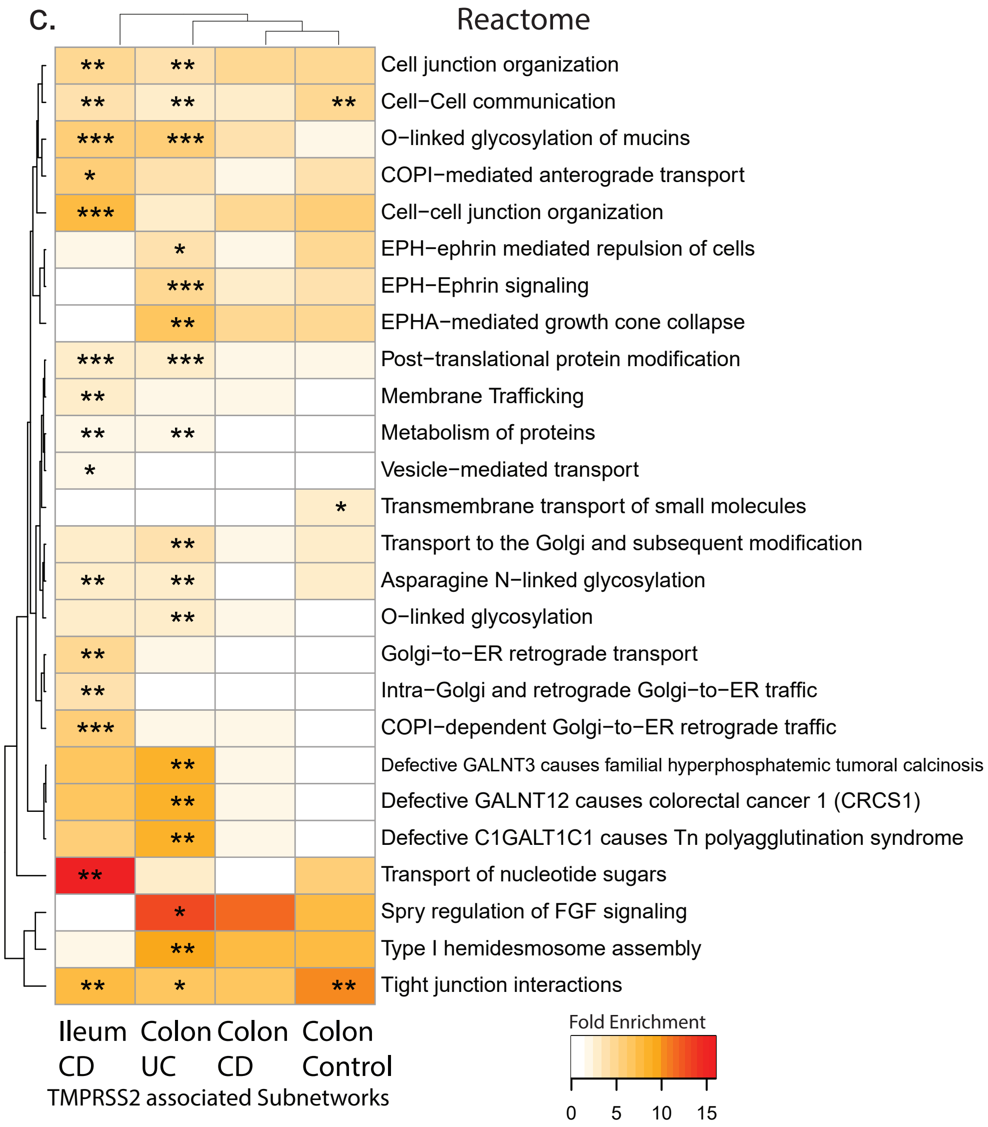

Adj P Value * = 0.1 -0.05, **=0.05-0.01, ***<0.01
